# Uninheritable but widespread bacterial symbiont mediates insecticide detoxification of an agricultural invasive pest *Spodoptera frugiperda*

**DOI:** 10.1101/2023.09.26.559648

**Authors:** Yunhua Zhang, Feng Ju

## Abstract

Identifying insecticide resistance-related insect bacterial symbiont is a vital compass for promoting symbiont-based insecticide resistance management and insecticide-free insect pest control. Known insecticide-resistance-related symbionts lack universality and genetic manipulation, as either not widespread or unculturable. Here, we discovered a widespread symbiont *Enterococcus casseliflavus* of a significant invasive insect pest, *Spodoptera frugiperda*. This symbiont enhances host insecticide resistance to chlorantraniliprole by amide bond breaking and dehalogenation-related insecticide degradation, suggesting its great potential for Lepidoptera pest insecticide resistance management. Complying with the increase in exposure risk of chlorantraniliprole, the *E. casseliflavus* of insects’ symbionts rather than mammals or environmental strains were notably enriched with putative chlorantraniliprole degradation genes. We also found that *E. casseliflavus* can transmit horizontally with high efficiency (100%) through cross-diet and cannibalism rather than vertical transmission from mother insect to offspring. Moreover, the widespread infection of *E. casseliflavus* in the field populations not only implies that an underlying symbiont-host co-evolution process driven by insecticide pressure might be underway, but also provides a novel therapeutic target of agricultural pests based on symbiont-targeted insect control (STIC).

## Introduction

Microbe-host interaction determines the adaptation and co-evolution of the holobiont, especially how it responds to the change of surrounding environment (Fischbach and Segre, 2016; Richardson, 2017; Li et al., 2023). In insect, gut microbiome is a large pool of functional microbial resources containing various active genes, especially those involved in nutrients synthesis and xenobiotic metabolism (Itoh et al., 2018; Manzano-Marı et al., 2020; Banerjee et al., 2022). By active gene expression, insect symbionts mediate a series of host phenotypes, including reproduction, development, behavior, immunity, and resistance to adversity (Moreira et al., 2009; Douglas, 2015; Liu and Guo, 2019). However, although the phenotypic effects of insect symbionts on hosts are common and stabilize in different insects, targeting specific functional symbionts at the strain level is particularly difficult due to the fact that the composition and diversity of microbiomes varies enormously within and between insect species (Kwong and Moran, 2016). Neither the causes nor the consequences of the mechanism of how the bacterium-provided active enzymes impact host phenotype are well understood, although many bacterial symbionts have been cultured and functionally verified in vitro (Pontes and Dale, 2006).

Insect symbionts have the detoxification potential for toxins, especially since the host lives under chronic stress from toxic substances (Kikuchi et al., 2012). In agriculture insect pests, insecticide is a joint toxic stress that dramatically affects insect holobiont, and there hence greatly enriches the insecticide-degrading symbionts in them, which may eventually lead to the development of insecticide resistance (Blanton et al., 2020). Insect hosts can obtain resistance against the insecticide fenitrothion by, for example, acquiring fenitrothion-degrading *Burkholderia* from the environment (Itoh et al., 2018). The phenomena are also noticed in a lab-selected *Bactrocera dorsalis* population, for which the gut symbiont *Citrobacter* sp. (strain CF-BD) plays a vital role in the trichlorphon degradation that contributes to increased trichlorphon resistance in *B. dorsalis* (Cheng et al., 2017). Although these studies indicated that bacterial symbionts in insects could mediate the host’s insecticide resistance by insecticide degradation, neither the insecticide degradation mechanism of bacterial symbionts nor how these symbionts infect and transmit within their host remain known.

The fall armyworm *Spodoptera frugiperda* has become a major global invasive pest in agriculture in the past decade. This crop pest, whether for native or invasive population, has developed insecticide resistance to pyrethroids, carbamates, diamides, *Bacillus thuringiensis* toxin, and organophosphates (Tay et al., 2023). Based on the variation in the insecticide application history at different geographical locations, the general nature of the evolution or development of insecticide resistance is still not systematically understood. Recently, microbiome composition variation of *S. frugiperda* is revealed to contribute to its diet-dependent insecticide susceptibility (Guo et al., 2022). Thus, we hypothesized that i) an invasive organism like an agricultural insect must have a strong gut for diet digestion, detoxification, and other essential acclimatization, and that ii) the gut microbiome is contributed to the insecticide resistance of insect pests, thus enhancing their field survival and invasive success across the globe. Here, we aim to 1) screen for and verify which pest insecticides can be detoxified and impacted by host bacterial symbionts; 2) identify how bacterial symbionts play a role in insecticide detoxification metabolism; and 3) resolve their field infection rate and transmission mechanism. The study will enrich the mechanistic understanding on the invasive success of a global crop pest facilitated by insecticide-degradation bacterial symbionts and create a novel green insecticide-free and green pest control solution based on the concept of symbiont-targeted insect control (STIC).

## Results

### Bacterial symbionts are involved in host’s insecticide susceptibility

Under the hypothesis on the host symbiont-facilitated insecticide detoxification, we chose six insecticides that commonly used in the field condition to check whether bacterial symbionts play a role in the insecticide detoxification of an agriculturally invasive pest *S. frugiperda* (Fig. 1a). The mixture of antibiotics (1000 mg/L of tetracycline and 1000 mg/L of rifampicin) was first used to inhibit bacterial symbionts, then, the susceptibility of *S. frugiperda* to each insecticide was assayed (Fig. 1b). The results showed that the bacterial symbiont load significantly dropped over 97% in the antibiotic-treated *S. frugiperda* compared to the control group (Fig. S1). After antibiotic treatment, the survival percentages of *S. frugiperda* in chlorantraniliprole, emamectin benzpate, chlorfenapyr, and lufenuron exposure were significantly decreased (*P* < 0.05, Fig. c-h). As for LC_50_ level, the susceptibility of *S. frugiperda* to chlorantraniliprole and chlorfenapyr were dramatically increased in antibiotic-treated *S. frugiperda* by 3.14, 1.29 and 2.59 times, respectively (Table 1). Notably, the LC_50_ of *S. frugiperda* to chlorantraniliprole was markedly dropped by approximately 3-fold (Table 1). Furthermore, the LC_50_ value of *S. frugiperda* to chlorfenapyr was also significantly decreased after antibiotics treatment. These results strongly indicated that bacterial symbionts contribute positively to the insecticide detoxification of their host *S. frugiperda* as a new invasive agricultural pest into China.

**Fig. 1.**
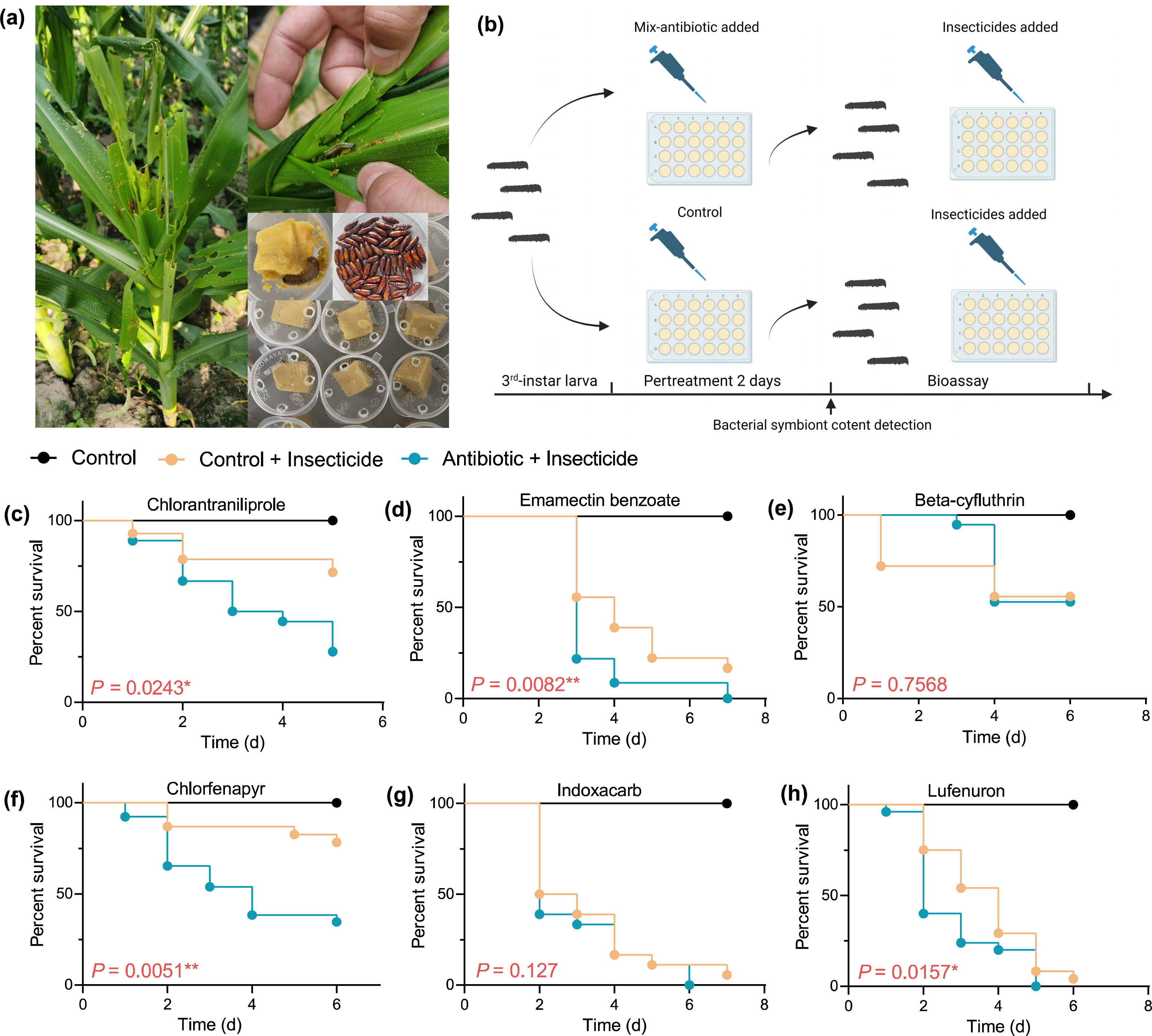
The influence of antibiotics on the susceptibility of *Spodoptera frugiperda* to six insecticides (in survival curve level). “*” and “**” represent significant difference between the Control and Antibiotics treatment groups in *P* < 0.05 and *P* < 0.01 level, respectively. The statistical analysis is based on the log-rank (Mantel–Cox) test.

**Table 1.** The influence of antibiotics on the susceptibility of *Spodoptera frugiperda* to insecticides.

| Insecticide | Treatment | N | Slope (SE) | LC<sub>50</sub> (95%CI) | χ<sup>2</sup> (df) | SR |
| --- | --- | --- | --- | --- | --- | --- |
| Chlorantraniliprole | Control | 144 | 1.19 (0.27) | 14.7 (7.70-60.86) | 4.25 (4) | **3.14\*** |
|  | Antibiotics | 144 | 2.04 (0.30) | 4.68 (3.51-6.83) | 4.30 (4) |  |
| Beta-cyfluthrin | Control | 144 | 2.07 (0.38) | 18.28 (10.26-25.72) | 2.71 (3) | 0.67 |
|  | Antibiotics | 144 | 3.12 (0.48) | 27.41 (20.37-34.26) | 3.05 (3) |  |
| Emamectin benzoate | Control | 144 | 2.55 (0.39) | 0.18 (0.13-0.23) | 7.23 (3) | 1.29 |
|  | Antibiotics | 144 | 3.24 (0.49) | 0.14 (0.1-0.17) | 5.41 (3) |  |
| Chlorfenapyr | Control | 144 | 5.14 (1.02) | 8.07 (5.25-10.16) | 1.98 (4) | **2.59\*** |
|  | Antibiotics | 144 | 3.68 (0.93) | 3.11 (0.67-5.36) | 0.80 (4) |  |
| Lufenuron | Control | 144 | 3.66 (0.65) | 0.57 (0.45-0.79) | 0.11 (3) | 1.14 |
|  | Antibiotics | 144 | 3.71 (0.63) | 0.50 (0.40-0.66) | 3.84 (3) |  |

Note: “\*” indicated that there was a significant difference in LC<sub>50</sub> between the antibiotic treatment group and the control group, and the analysis was based on Polo Plus; N:Insect number; SR (Susceptibility Ratio) = Control (LC<sub>50</sub>)/ Antibiotics (LC<sub>50</sub>).

### The response pattern of bacterial symbionts on insecticide exposure

Once bacterial symbionts are demonstrated to involve in insecticide detoxification of invasive pest *S. frugiperda*, we next explored the response pattern of bacterial symbionts under insecticide exposure to anchor the core bacterial symbiont taxa involved in host insecticide detoxification (Fig. S2a). By 16S rRNA gene amplicon sequencing, we found that Firmicutes were the core and predominant bacterial symbiont of *S. frugiperda* (relative abundance > 90%, Fig. S2b). The β-diversity of bacterial symbiont was significantly changed in response to the exposure of chlorantraniliprole, but not other insecticides (Fig. 2a). The α-diversity indexes, including observed species (or ASVs) and Shannon index, were significantly increased during chlorantraniliprole exposure compared to the control (Fig. 2b and c). The increasing microbiota α-diversity under chlorantraniliprole exposure was reflected by the increase in relative abundance of bacterial symbiont species, including *Lactobacillus plantarum*, *Pseudomonas* sp., *Enterococcus casseliflavus* and *Lactobacillus brevis* (Fig. 2d and e). However, since the bacterial symbionts of *S. frugiperda* are predominated by *Enterococcus* (90.0% to 99.1%), we postulated that the *Enterococcus* symbionts, which were enriched significantly (e.g., *E. casseliflavus* increased by 10.81 times, *P* = 0.034) after chlorantraniliprole stress, should be involved in the detoxification of the host to chlorantraniliprole.

**Fig. 2.**
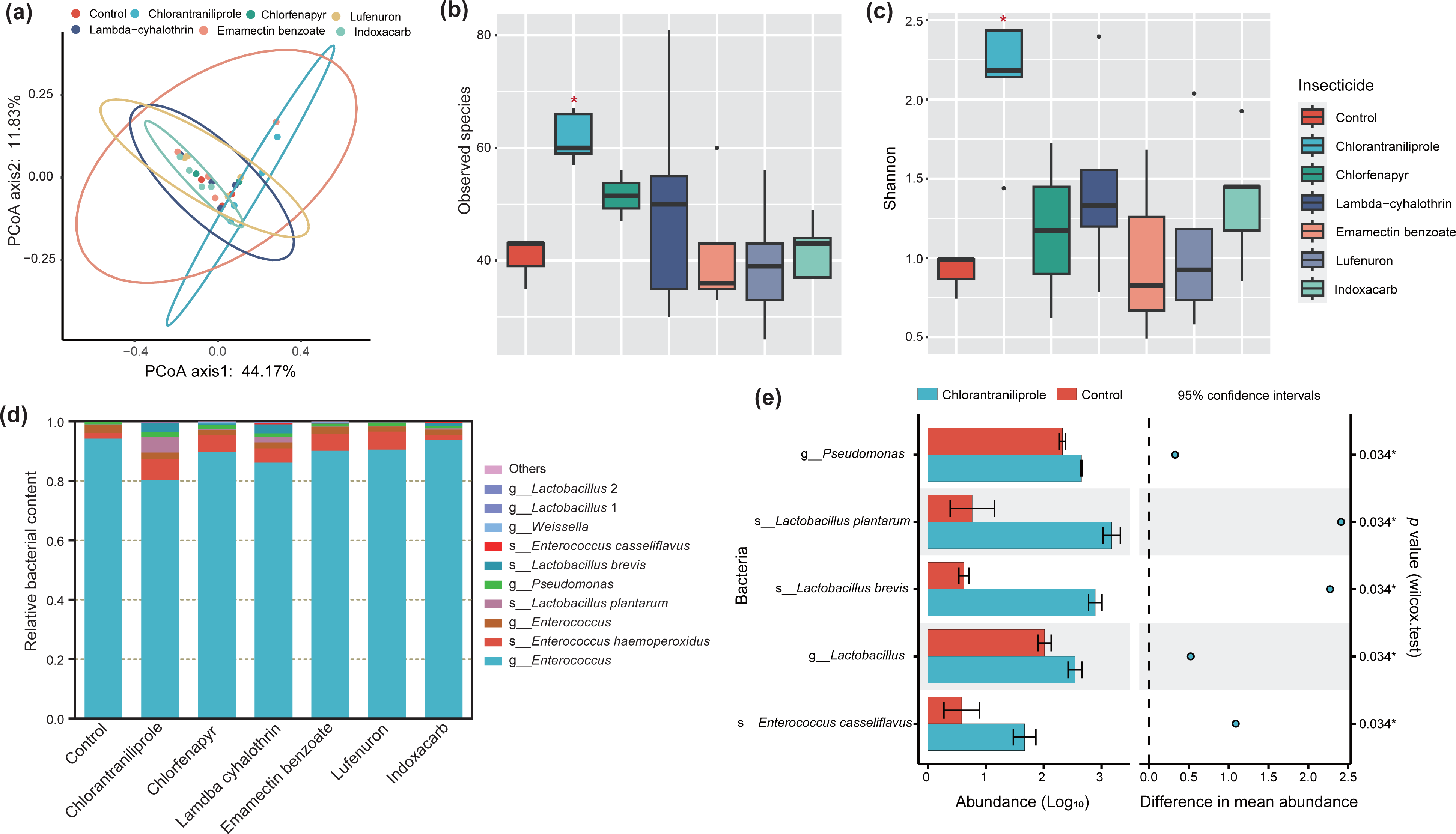
Response pattern of bacterial symbionts of *Spodoptera frugiperda* under the different insecticide’s exposure. **(a)**: PCoA of *S. frugiperda* microbiome under the different insecticide’s exposure; **(b and c):** α-diversity of *S. frugiperda* microbiome under the different insecticide’s exposure; **(d):** Taxonomic composition of *S. frugiperda* microbiome species assignable at the species (s) or only genus (g) level under the exposure to different insecticides; **(e):** Differential species abundance profiles of bacterial symbionts between the control group and the chlorantraniliprole exposure group; “*” represent significant difference between treatment and control group (*P* < 0.05), statistical analysis is based on the Wilcox-test.

### A bacterial symbiont of *Enterococcus* mediates host’s chlorantraniliprole susceptibility by insecticide degradation

To ascertain the core bacterial symbiont strains involved in host’s chlorantraniliprole detoxification, we obtained 4 bacterial isolates from the larvae gut of *S. frugiperda* pre-exposed with chlorantraniliprole (Fig. 3a) and further taxonomically identified them by full-length 16S rRNA gene sequences as *E. casseliflavus*, *Enterococcus mundtii*, *Klebsiella variicola* (NCBI accession ID: MZ475068) and *Klebsiella oxytoca*, respectively (Fig. S3). We hypothesized that bacterial symbionts may mediate host’s detoxification by degrading insecticide to the innoxious or less toxic substances. To this end, the chlorantraniliprole degrading ability of these four bacterial strains were monitored by ultraviolet spectrophotometer. The results showed that a new isolate of bacterial strain, which first named here as *E. casseliflavus* EMBL-3, effectively degraded chlorantraniliprole in 3 days by 25.9% (from 20.0 mg/L to 14.8 mg/L). In contrast, the other three bacterial isolates did not significantly degrade chlorantraniliprole (*P* > 0.05) (Fig. 3b). To verify this result, we then measured chlorantraniliprole degradation effectively via LC-MS/MS that confirm *E. casseliflavus* degraded 24.2% of chlorantraniliprole in 3 days (Fig. S4 and 3c). We further analyzed the impact of *E. casseliflavus* on the insecticide susceptibility of *S. frugiperda* (Fig. S5). We discovered that the susceptibility of *S. frugiperda* to chlorantraniliprole was dramatically decreased (by 4.7-fold) with *E. casseliflavus* feeding, while another bacteria isolate *K. variicola* which cannot degrade chlorantraniliprole in vitro showed no significant effect on host chlorantraniliprole susceptibility (*P* = 0.82, Fig. 3d and Table S1). These results suggested that *E. casseliflavus* symbionts significantly reduce their host’s chlorantraniliprole susceptibility by insecticide biodegradation.

**Fig. 3.**
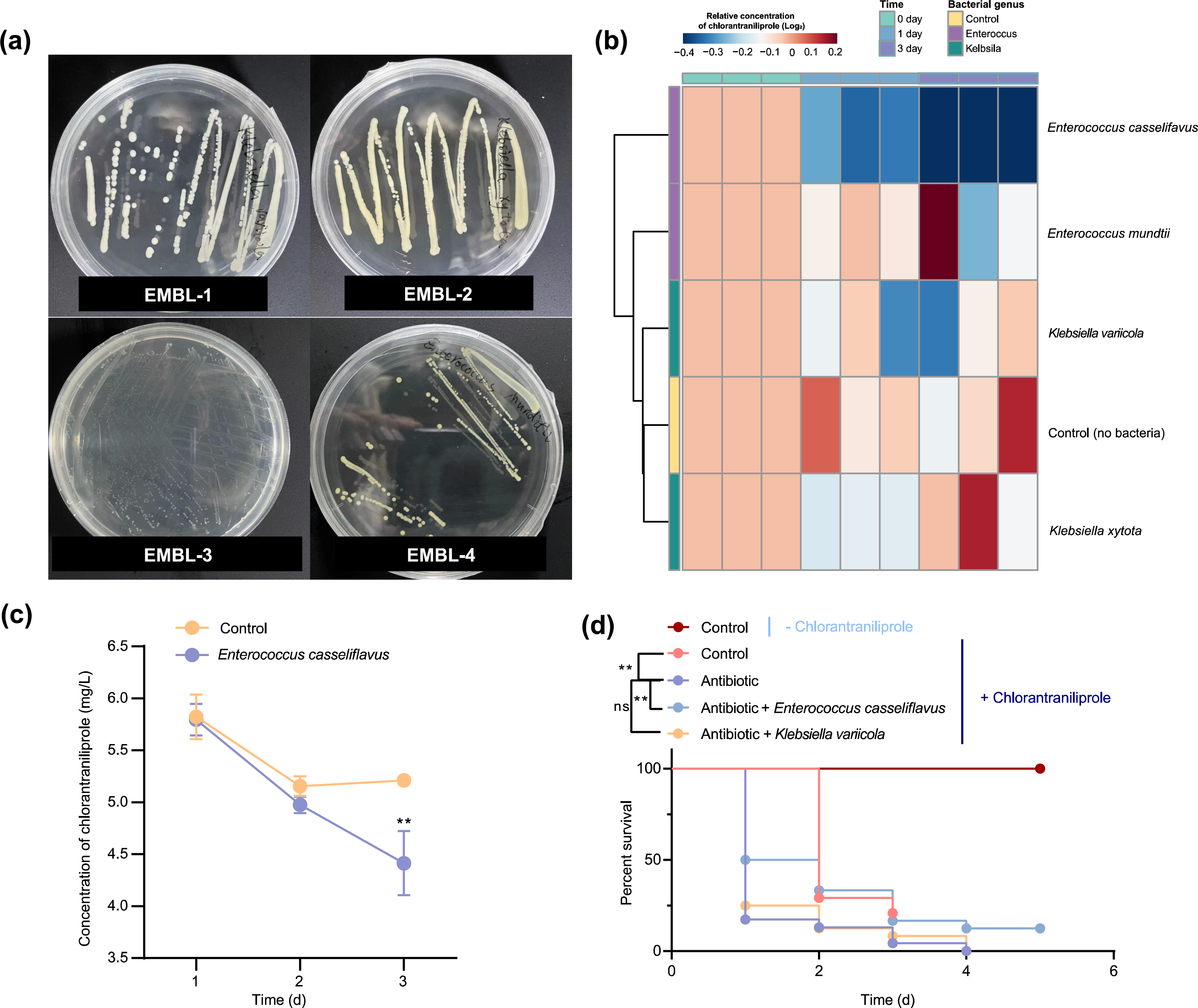
A bacterial isolate *Enterococcus casselifiavus* increased host’s insecticide detoxification by chlorantraniliprole degradation. **(a)**: Four bacterial isolates from *Spodoptera frugiperda*; **(b):** Chlorantraniliprole degrading ability of four bacterial isolates. Heatmap shows the relative content of chlorantraniliprole after bacterial symbionts treatment; **(c):** Degradation effective of *E. casselifiavus* on the chlorantraniliprole based on the LC-MS method; **(d):** The influence of *E. casselifiavus* and *Klebsiella variicola* on chlorantraniliprole susceptibility of *S. frugiperda.* “**” represent significantly difference between control and treatment group (*P* < 0.01). The statistical analysis is based on the log-rank (Mantel–Cox) test.

### Symbiont-facilitated insecticide detoxification mechanism indicates holobiont and co-evolution under insecticide selection

To further explore the degradation mechanism of *E. casseliflavus* on chlorantraniliprole, we performed two-part protein degradation activity assays to distinguish the intracellular and extracellular biodegradation process (Fig. S6a). The extracellular and intracellular proteins were first extracted from *E. casseliflavus* fermentation broth and cells, respectively. By co-incubation chlorantraniliprole with extracellular and intracellular proteins for 3 days, we then found that extracellular proteins exhibited 5.16 times higher chlorantraniliprole-degrading efficiency than intracellular proteins (Fig. 4a). Specifically, the concentration of chlorantraniliprole (mg/L) was decreased by 18.6% post treatment with *E. casseliflavus* extracellular protein (EEP), whereas less than 1% decrease post treatment with the *E. casseliflavus* intracellular protein (EIP, Fig. 4a). Moreover, the protein expression pattern was changed dramatically in extracellular proteome much more than the intracellular proteome (Fig. S6b). In the extracellular proteome, we identified 543 differentially expressed proteins, including 260 over-expressed and 283 down-regulated proteins (Fig. 4b). In contrast, only 30 and 44 proteins showed over-expression or down-regulated expression in intracellular protein between the control and the chlorantraniliprole treatment, respectively (Fig. 4c). Interestingly, we discovered the intracellular over-expression of secretion system and protein export system proteins in chlorantraniliprole-treated *E. casseliflavus* cells which strongly implied that *E. casseliflavus* degradation of chlorantraniliprole was largely dependent on the extracellular protein (Fig. 4c, Table S2).

**Fig. 4.**
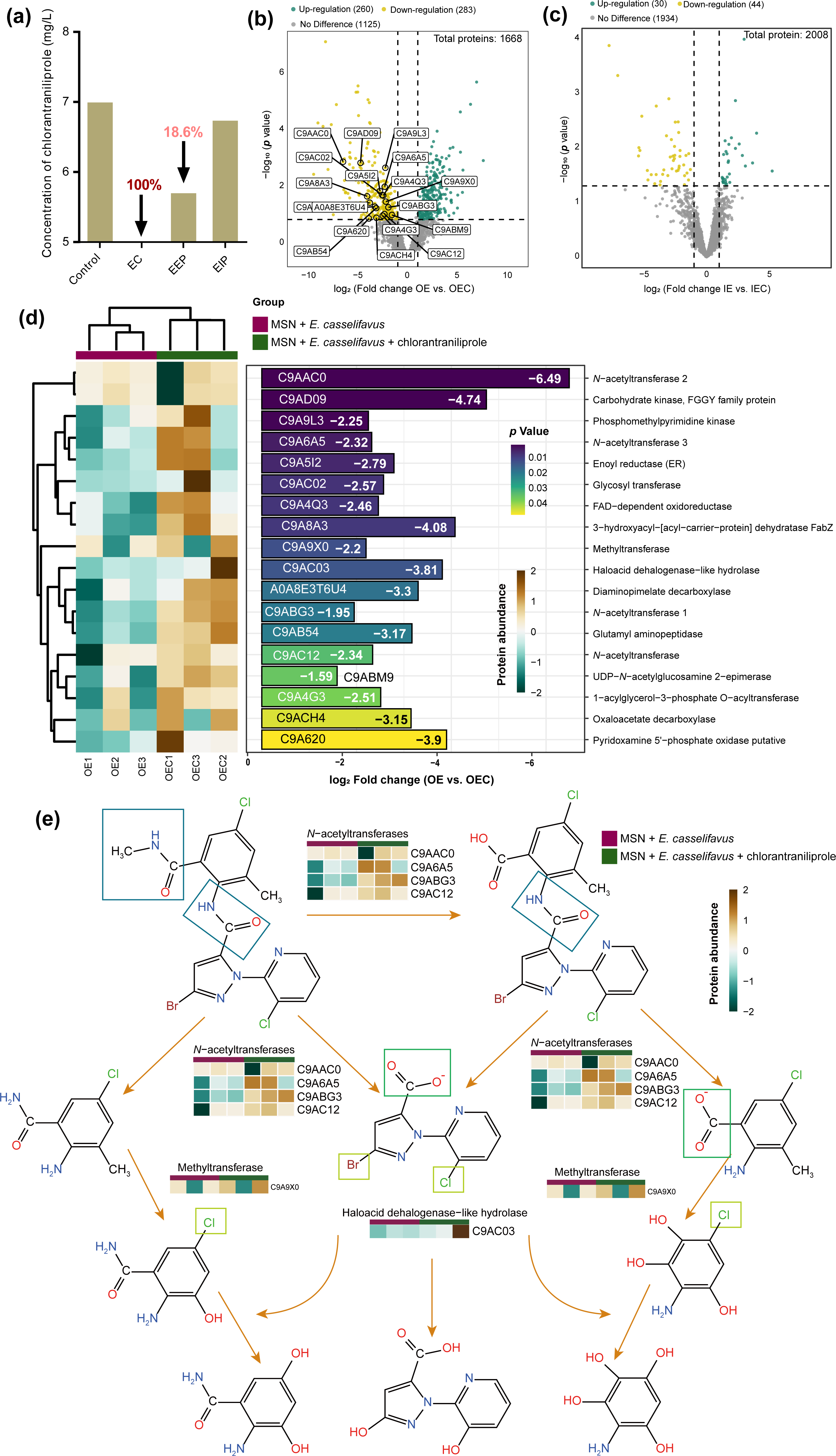
Degradation mechanism of *Enterococcus casselifiavus* on chlorantraniliprole. **(a)**: Degradation efficiency of *E. casselifiavus* and its intracellular and extracellular proteins to chlorantraniliprole. Control: MSM + chlorantraniliprole; EC: MSM + *E. casselifiavus* + chlorantraniliprole; OEP: MSM + chlorantraniliprole + extracellular proteins; IEP: MSM + chlorantraniliprole + intracellular protein; **(b and c)**: Volcano Plot of different expression protein after chlorantraniliprole exposure in intracellular (c) and extracellular (b) proteins. (b) highlight 18 proteins that significantly over-expression in extracellular proteins after chlorantraniliprole exposure and have more possible chlorantraniliprole degrading potential. OE: Extracellular proteins expression level in MSM + *E. casselifiavus*; OEC: Extracellular proteins expression level in MSM + *E. casselifiavus* + chlorantraniliprole; IE: Intracellular proteins expression level in MSM + *E. casselifiavus*; IEC: Intracellular proteins expression level in MSM + *E. casselifiavus* + chlorantraniliprole. The significant level set as *P* < 0.05, and the green and yellow point shows significantly down or over-expression protein, respectively (Fold change > 2); **(d)**: Abundance, fold change and the *P* value of highlight proteins that have more possible chlorantraniliprole degrading potential. **(e):** Putative chlorantraniliprole degradation pathway of *E. casselifiavus*. The pathway was proposed based on multi-omic analyses that integrated the genomic, proteomic results for *E. casselifiavus* during growth on MSN containing chlorantraniliprole. Dehalogenation and breakdown of amide bonds are the dominant mechanism of *E. casselifiavus* degradation of chlorantraniliprole.

Among the over-expressed extracellular proteins, we screened for putative chlorantraniliprole-degrading enzymes (fold change > 2) according to the function annotation information of proteins and combined with the compound structure of chlorantraniliprole (Fig. 4d), such as *N*-acetyltransferase (3.86-89.88), carbohydrate kinase (26.72), glycosyl transferase (5.93), FAD−dependent oxidoreductase (5.50), methyltransferase (4.59), and haloacid dehalogenase (14.03). Especially, considering the fact that chlorantraniliprole contains 2 amide and 3 halogen bonds, the identification of 4 *N*-acetyltransferases and 1 dehalogenase implied that *E. casseliflavus* can degrade chlorantraniliprole via dehalogenation and amide bond break (Fig. 4d). According to the biodegradation products detected via LC-MS/MS (Fig. S7), and the biodegradation pathway predicted by Biocatalysis/Biodegradation Database of (EAWAG-BBD) (Fig. S8, Sivakumar et al., 2017), we propose the first hypothetical biodegradation pathway of chlorantraniliprole by *E. casseliflavus* (Fig. 4e), a general insect symbiont of Lepidoptera insects (Gomes et al., 2023).

To further deduce the co-evolution significance of chlorantraniliprole-degrading enzyme in *E. casseliflavus*, it is hypothesized that the ability of bacterial symbionts to evolve insecticide biodegradation is the consequence of host-microbe co-evolution driven by insecticide stress (e.g., widespread in agricultural fields). To test the hypothesis, we sequenced and constructed the whole genome sequence of *E. casseliflavus* and excitingly discovered this symbiont harbored plentiful putative chlorantraniliprole-degrading genes (Fig. S9), including 32 acetyltransferases, 8 dehalogenase and 19 methyltransferase genes. Echoing this intriguing finding, we further identified that an *E. casseliflavus* isolate from *Bombyx mori* also harbors highly abundant putative chlorantraniliprole-degrading genes (39 acetyltransferases, 6 dehalogenase and 20 methyltransferase genes), whereas *E. casseliflavus* isolates from environment (e.g., soil and wastewater) and mammal gut (e.g., human baby and hog) contain much less (< 5) or almost no putative chlorantraniliprole-degrading genes (Fig. 5). We also found 6 acetyltransferases (4 in *E. casseliflavus* of *B. mori* and 2 in *E. casseliflavus* of *S. frugiperda*) and 1 dehalogenase (in *E. casseliflavus* of *S. frugiperda*) in the unique gene cluster of insect’s *E. casseliflavus* (Fig. 5). Our discovery of the dramatic enrichment of chlorantraniliprole-degrading genes in insect symbionts genomes as well as the prominent differentiation in biotope variation consisted with the well-acknowledged insecticide exposure risk in the following decreasing order: insect > environment > mammal. The above findings together suggested for the first time that symbiont-host interaction shaped by insecticide biodegradation promotes invasive pest holobiont (i.e., *S. frugiperda* and their symbionts) evolution to resist insecticide pressure in agriculture.

**Fig. 5.**
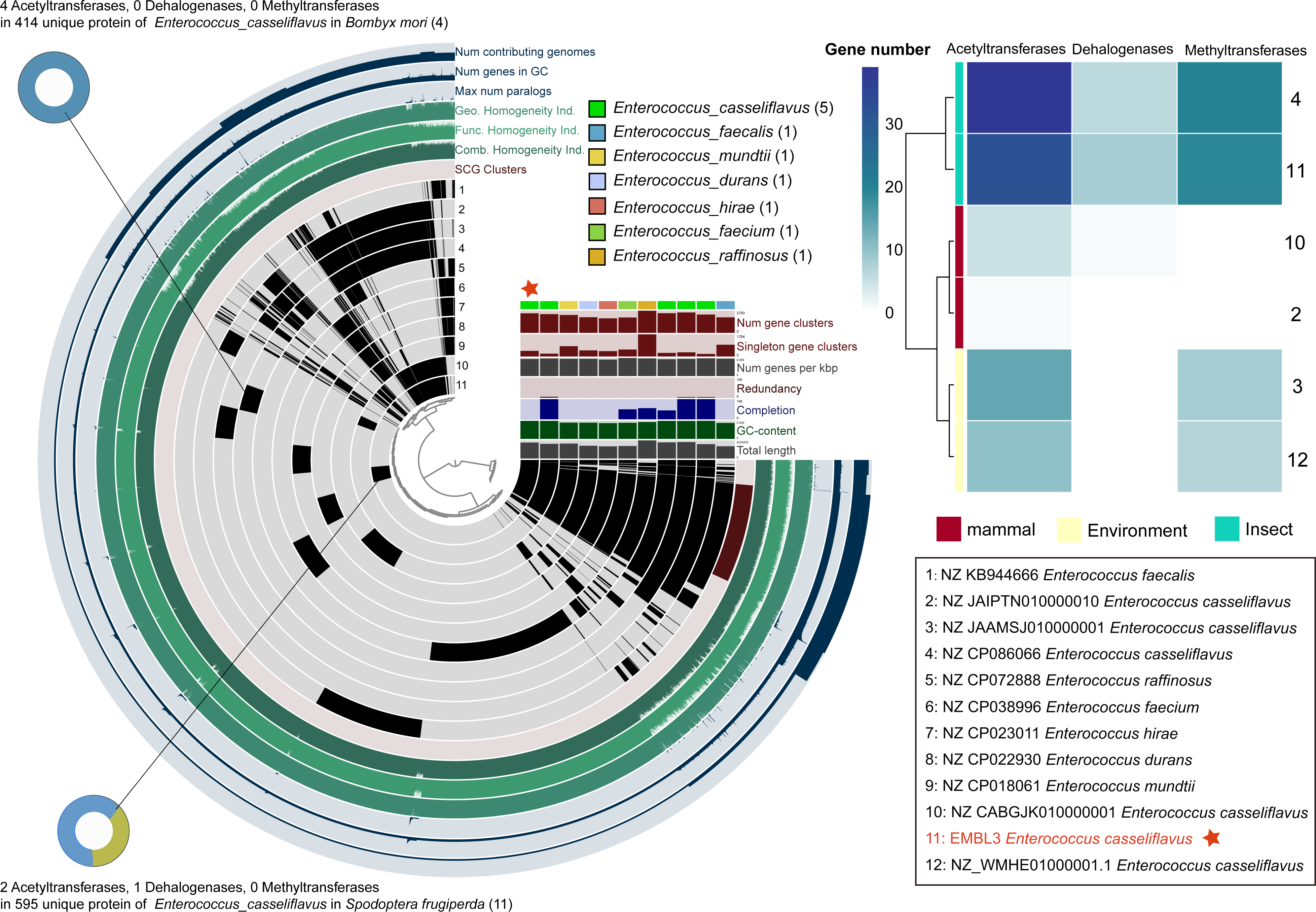
Pangenomic analysis and potential chlorantraniliprole degrading gene distribution of *Enterococcus* genomes by Anvi’o. The central plot of the interface represents a hierarchical clustering dendrogram based on gene presence/absence. In the circle interface, each layer (grey) represents all genes (black) in a single genome. The unique gene cluster of each genome were identified by BPGA, then annotation with eggnog-mapper, All and specific potential degradation genes of chlorantraniliprole were counted and show in heatmap and sector diagram.

### Unherdable but widespread *E. casseliflavus* symbionts with efficient horizontal transmission ability

To tap the application prospect of *E. casseliflavus* symbionts in insecticide resistance management and insect pest control, we are particularly interested in unraveling the infection frequency of *E. casseliflavus* in the agricultural field as well its spread mechanism within and between insect populations. Because widespread infection and easy transmission are the main factors limiting the application of targeted symbionts in insect control, we collected five field populations of *S. frugiperda* across China to monitor infection frequency of *E. casseliflavus* (Fig. 6a). The results showed that all *S. frugiperda* population individuals infected *E. casseliflavus* with 100% frequency (Fig. 6a). Driven by our first discovery that infection of *E. casseliflavus* symbionts is widespread in *S. frugiperda* field populations, the vertical propagation ability of *E. casseliflavus* in host’s different generations was further examined (Fig. 6b). The special primer set ECHP-F and ECHP-R was designed to target a unique hypothetical protein-coding gene (NCBI accession no.: WP_260661680.1) of *E. casseliflavus* compared to other *Enterococcus* which was bioinformatically found via pangenomic analysis (Fig. 6b). The experimental result demonstrated that *E. casseliflavus* could be fed back into the host and continued to proliferate until the pupal stage after inoculation was ceased, but the seeding effect was distinct between the male and female adult worms. The males had a partial symbiotic bacterial load, while the females had a very low and negligible abundance leading to the loss of infection from the eggs to larvae of the next generation (Fig. 6b). It is further speculated that maintaining a high population infection frequency may be realized through horizontal transmission rather than vertical transmission from mother to offspring insect. In this case, the horizontal transmission of *E. casseliflavus* between individual *S. frugiperda* was further assessed. Two relevant scenarios are considered: cross-feeding and cannibalism (Fig. 6c). The results revealed that *E. casseliflavus* can efficient transmit horizontally via both food crosses and cannibalism (Fig. 6c and d), suggested that horizontal transmission is a key route to maintaining the widespread infection of *E. casseliflavus* bacterial symbionts in field *S. frugiperda* population.

**Fig. 6.**
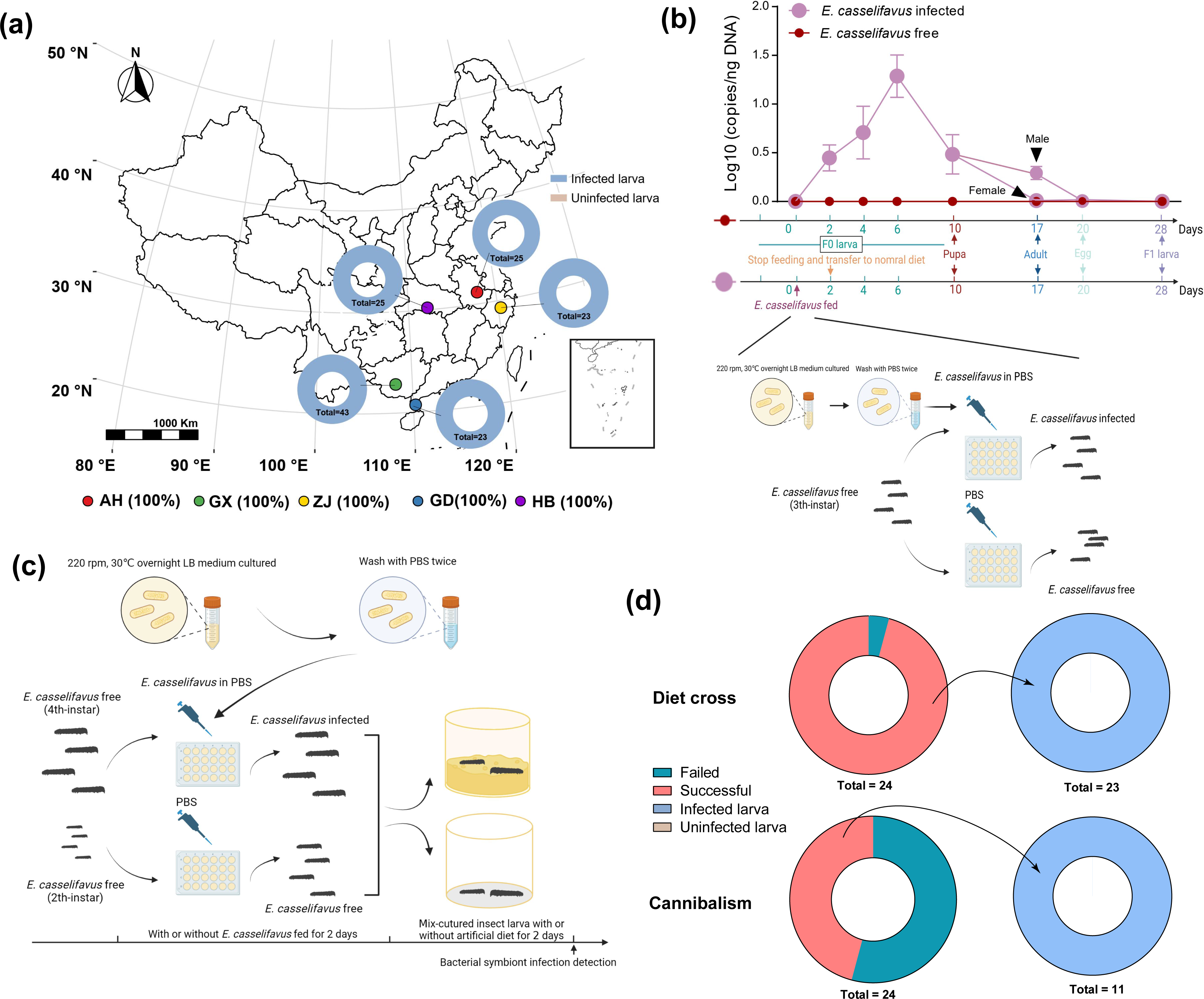
Field infection rate and transmission pattern of *Enterococcus casselifiavus* in *Spodoptera frugiperda.* **(a)**: Infection rate of *E. casselifiavus* in 5 field populations of *S. frugiperda*. **(b):** Infection pattern of *E. casselifiavus* in laboratory population from F0 to F1 generation. **(c):** Diagram of horizontal transmission capability detection. **(d):** Horizontal infection efficiency.

## Discussion

To combat with adversity, insect and its symbionts as a holobiont have been co-evolving to encode various nutrient synthesis and xenobiotic detoxification functional genes in their hologenome (Pester and Brune, 2007; Rosenberg and Zilber-Rosenberg, 2018; Zhang et al., 2021). The current study identified a widespread infection of bacterial symbiont in field host populations, i.e., *E. casseliflavus*, which is first proved to enhance host insecticide resistance to chlorantraniliprole by insecticide degradation via dehalogenation and amide bond break pathway in an agriculturally invasive, insecticide resistant insect pest, *S. frugiperda*. Our discovery revealed that beneficial and universal mutualism between *S. frugiperda* and *E. casseliflavus* may contributed to the global invasive success of *S. frugiperda*. Moreover, our firsthand results highlight a progress of symbiont-host co-evolution driven by insecticide pressure is faster than priorly appreciated, and support *E. casseliflavus* symbionts as a suitable and new bacterial target for insecticide-resistant Lepidoptera pest control.

To identify the bacterial symbiont involved in the insecticide response of *S. frugiperda*, we found that chlorantraniliprole exposure significantly changed host gut bacterial community diversity, and considerably enriched for 5 bacterial species (Fig. 2), of which *E. casseliflavus* strain EMBL-3 was isolated and functionally verified as chlorantraniliprole-detoxifying symbionts both in vivo and in vitro. To our knowledge, this is the first report of *E. casseliflavus* as an insecticide detoxification and resistance mediating bacterium in *S. frugiperda*, a major invasive pest in the world. Consolidating our finding on *Enterococcus* symbiont-facilitated insecticide resistance via biodegradation, other different bacterial symbionts (e.g., *Pseudomonas* and *Lactobacillus*) were noticed to be enriched in chlorantraniliprole-exposed *S. frugiperda*. Interestingly, a *Pseudomonas* strain isolated from chlorantraniliprole waste soil can also degrade chlorantraniliprole (Almeida et al., 2017; Gao et al., 2019). In addition, the chlorfenapyr susceptibility of *S. frugiperda* was also dramatically changed which implied that the bacterial symbiont can involve in chlorfenapyr degradation but may be different strain. The above results suggested that *S. frugiperda* gut symbionts are an excellent reservoir of insecticide-degrading bacteria with bioremediation potential.

The key ecological question of *S. frugiperda* gut colonization by insecticide-degrading bacterial symbionts is where these insecticide-degrading bacteria come from. In the absence of experimental support, previous researches revealed that caterpillar lacked a resident gut microbiome whose diversity was dependent on the soil rather than plant host (Hammer et al., 2017; Hannula et al., 2019). Thus, we hypothesize that the source of degrading bacterial symbionts be from soil (Huang et al., 2018). Numerous insecticide residue leads to the enrichment of numerous insecticide-degrading bacteria in farmland soil, which is beneficial in environmental remediation and pollutants transformation (Fenner et al., 2013). However, unfortunately, when these bacteria are transferred to insect pests through plants, the insect may rapidly resist to the corresponding insecticides, leading to the failure of pest control and serious crop loss in agriculture (Tago et al., 2015; Itoh et al., 2018). Although the influence of insecticide resistance evolution of insect infection with insecticide-degrading bacterial symbionts remains unknown, the similar results from previous studies indicated that insects containing insecticide-degrading bacteria such as *Riptortus pedestris*, *Cletus punctiger*, *Bactrocera dorsalis* and *Aphis gossypii*, could evolve more quickly in the insecticide resistance than was priorly thought before under the help of bacterial symbionts (Cheng et al., 2018; Sato et al., 2021; Ishigami et al., 2021; Lv et al., 2023). Furthermore, our first evidence for extensive infection of these insecticide-degrading bacterial symbionts in field strains *S. frugiperda* implies that the potential for insecticide resistance evolution in field insect pests is much greater than previously realized.

Based on comparative genomic and pan-genomic analyses to expound the particularity of chlorantraniliprole-degrading symbionts, we found that the potential chlorantraniliprole degradation genes were notably enriched in insect symbiont genomes rather than those symbionts of mammals or environment bacterial strains, and increased with the chlorantraniliprole exposure risk. Lepidopterans are shown to have the highest average number of horizontal gene transfer (HGT)-acquired genes from bacteria (Li et al., 2021). In a perspective, potential insecticide-degrading genes are prominently enriched in insect bacterial symbionts, implying that these xenobiotic-degrading genes are likely to be integrated into the host genome by plasmid or transposon to form a horizontal transfer of insecticide resistance genes (Acuña et al., 2012). However, compared to the evolutionary history of insects, insecticide use should be far from long enough to trigger direct selection of host resistance, but this assumption does not negate either the occurrence of HGT in future long-term and high-stress pesticide application scenarios, or the likelihood in co-selection by ancient environmental stress (such as plant secondary metabolites). Of course, when host insects acquire insecticide-degrading genes, it also means that their symbiotic relationship with bacterial symbionts may be broken down.

Besides our discovery of *E. casseliflavus* symbionts-facilitated host pest detoxification and holobiont evolution, we also found that these insecticide-degrading bacterial symbionts cannot transmit vertically from mother to offspring of *S. frugiperda*. This novel finding agreed with previous studies that indoor insect populations had lower symbiotic bacterial diversity, implying that the symbionts can be lost in indoor long time artificial diet feeding (Kakumanu et al., 2018; Zhang et al., 2018; Zheng et al., 2023). Exploring pest resistance mechanisms in artificial diet mode may be deficient, especially when some artificial feeds are supplemented with numerous antibiotics, accelerating the decline of symbionts after field population transfer to indoor feeding. From this perspective, what happens in the artificial diet context of this and prior laboratory studies cannot completely replace what happens in the field (Pang et al., 2018; Zhang et al., 2023). Therefore, future studies on corn-feeding or wide-type *S. frugiperda* are needed to complement and further validate the original mechanism understanding of bacterial symbiont-mediated host detoxification achieved here.

In conclusion, our study discovered *E. casseliflavus* as the uninheritable but widespread gut bacterial symbionts of *S. frugiperda* that mediate host chlorantraniliprole detoxification by insecticide degradation. The widespread infection rate in the field populations and simple transmission pattern within population make *E. casseliflavus* to have great applications potential for symbiont-targeted insect pest control and environmental bioremediation of chlorantraniliprole contamination. Our new findings together provide a new perspective for understanding the native adaptation and success rate of a globally emerging agricultural invasive insect pest, our new findings laying the foundation for the future of symbiont-targeted agricultural injurious insect suppression (STIS) or economic insect promotion (STEP).

## Materials and Methods

### Insects and chemicals

*S. frugiperda* used in this study was originally collected in a maize field in Nanning, Guangxi of China (108.28 °E, 23.16 °N) in September 2022. The field populations of *S. frugiperda* were also collected in five provinces of China, including Zhejiang, Guangdong, Anhui, Hubei and Guangxi at 2022 (Table S3). The larvae were fed on an artificial diet according to a previous study (Bolzan et al., 2019). Adults were fed in 30 cm× 30 cm× 30 cm insect cages with 10% honey solution to nutritional supplements. All insects were reared at 25 ± 1 °C and 65 ± 5% relative humidity (RH) with a light:dark (L:D) photoperiod of 16 h:8 h.

Information on the chemical insecticides and antibiotics used in this study is listed in Table S4. Stock solutions of the insecticides were diluted with acetone or *N*, *N*-dimethyl formamide to the desired concentrations.

### Antibiotic treatment and insecticide bioassay

The diet-overlay method was used for insecticide bioassays as described previously with a slight modification (Bolzan et al., 2019; Guo et al., 2022). In brief, 900-μL artificial diet was added in a well belonging to 24-well plates, after cure, 100 μL of the insecticide solution was added to the diet surface and ensuring even coverage. Five concentrations of insecticide (prepared diluent with distilled water containing 0.1% Triton X-100) and control (0.1% Triton X-100 distilled water) were performed to bioassay for *S. frugiperda*. After the insecticide solution dried, 24 healthy 3rd-instar larvae of *S. frugiperda* were carefully individually placed in each well of plates representing three replicates in each concentration. The treated insect larvae were kept in the insect-rearing environment and the mortality was recorded at 24 h (lambda-cyhalothrin and chlorfenapyr), 48 h (indoxacarb, emamectin benzoate and chlorantraniliprole) after treatment. The survival probole analysis was performed consistent with the previously described using an appropriate sublethal concentration of insecticide (LC_50_), and the survival was checked every 24 h after being treated with insecticide.

Antibiotic treatment was consistent with the previous study by the diet-overlay method, 100 μL of the mixed antibiotic solution (1000 mg/L of tetracycline and 1000 mg/L of rifampicin) was added to the surface of the artificial diet in each well, ensuring even coverage of the diet surface evenly. Third-instar larvae were transferred to the antibiotic-covered diet for 24 h for pretreatment with antibiotics (Guo et al., 2022). The pretreated larvae were used for further bioassay experiments.

### DNA extraction and 16S rRNA amplicon sequencing

Total genomic DNA of indoor and field populations *S. frugiperda* was extracted using the OMEGA Tissue DNA Kit (Omega Bio-Tek, Norcross, GA, USA), following the manufacturer’s instructions. Extracted DNAs were quantity and quality measured using a NanoDrop NC2000 spectrophotometer (Thermo Fisher Scientific, Waltham, MA, USA) and agarose gel electrophoresis, respectively.

PCR amplification of the bacterial 16S rRNA gene V3–V4 region was performed using the forward primer 338F (5’-ACTCCTACGGGAGGCAGCA-3’) and the reverse primer 806R (5’-GGACTACHVGGGTWTCTAAT-3’). Sample-specific 7-bp barcodes were incorporated into the primers for multiplex sequencing. The PCR components contained 5 μL of buffer (5×), 0.25 μL of Fast pfu DNA Polymerase (5 U/μL), 2 μL (2.5 mM) of dNTPs, 1 μL (10 uM) of each Forward and Reverse primer, 1 μL of DNA Template, and 14.75 μL of ddH_2_O. Thermal cycling consisted of initial denaturation at 98 °C for 5 min, followed by 25 cycles consisting of denaturation at 98°C for 30 s, annealing at 53°C for 30 s, and extension at 72°C for 45 s, with a final extension of 5 min at 72°C. PCR amplicons were purified with Vazyme VAHTSTM DNA Clean Beads (Vazyme, Nanjing, China) and quantified using the Quant-iT PicoGreen dsDNA Assay Kit (Invitrogen, Carlsbad, CA, USA). After the individual quantification step, amplicons were pooled in equal amounts, and pair-end 2×250 bp sequencing was performed using the Illlumina NovaSeq platform with NovaSeq 6000 SP Reagent Kit (500 cycles) at Shanghai Personal Biotechnology Co., Ltd (Shanghai, China).

### Enrichment, isolation, and identification of chlorantraniliprole-degrading bacterial symbiont strain

Based on our results, chlorantraniliprole is the most fold-change in susceptibility after antibiotics treatment, and also the most host microbiome responsive insecticide, so we choose chlorantraniliprole for the next study to explain the mechanism of microbiome mediated insecticide detoxification. To isolate chlorantraniliprole-degrading bacterial symbionts, the gut of ten *S. frugiperda* larvae pre-exposed to chlorantraniliprole were suspended in 10 mL of PBS and vortexed for 5 min eluting symbiotic bacterial cells. Microcentrifugation (approximately 500 rpm) removes the host’s solid tissue and gut contents. Then the solution was gradient diluted by PBS by 10× from 10^-1^ to 10^-8^ and 50 μL diluent solution was evenly coated on the LB plate media without antibiotics and cultured overnight at 30°C. The next day, different single colonies were selected and added into the 500 μL LB liquid medium without any antibiotics for expanded culture, and plate streak purification and species identification were carried out.

For bacterial symbiont strains identification, a near full-length 16[S rRNA gene sequence was PCR amplified using the universal primers 27[F (5’-AGAGTTTGATCCTGGCTCAG-3’) and 1492[R (5’-GGTTACCTTGTTACGACTT-3’).

The 16⎕S rRNA gene amplicon sequence obtained from Sanger sequencing was deposited in the National Center for Biotechnology Information (NCBI) database and annotated using NCBI’s online Basic Local Alignment Search Tool (BLAST) on 20 September in 2022 based on the BLAST+ 2.13.0. The phylogenetic trees were constructed via MEGA 11 trough Maximum likelihood (ML) method with 1000 Bootstraps replications.

### Biodegradation of chlorantraniliprole by bacterial symbiont isolates of *S. frugiperda*

The obtained bacterial symbiont isolates of *S. frugiperda* were used to measure the biodegradation ability of chlorantraniliprole according to a previous study with slight modification (Gao et al., 2019). Briefly, the isolate strains were incubated in LB media at 30°C and 220 rpm until the OD_600_ of the culture solution reached 1.0. Then, the strain was aseptically collected (5000 rpm, 5 min), washed twice with PBS solution, and resuspended in an equal volume of PBS solution. To test the degradation ability of isolate strains, 10% of resting cells (OD_600_ = 1.0) was added to a 20 mL minimal salt liquid medium (MSM) containing chlorantraniliprole (6 mg/L). It was incubated at 30°C and 220 rpm. The non-inoculated culture served as a control.

The residual concentration was extracted by ethyl acetate and measured by ultraviolet spectrophotometer at 265 nm. After preliminary screening, the strains with the most chlorantraniliprole-degrading potential are screened for subsequent precise experiments. After extraction by ethyl acetate, the solution was dried with nitrogen and redissolved with 80% methanol then measured by HPLC following method: the separation was carried out on theHSS T3 column (100 mm × 2.1 mm, 1.8 μm) at 40°C. The mobile phase A was Ultra-pure water containing 0.1% FA and the mobile phase B was methanol containing 0.1% FA. The flow rate was 0.3 mL/min, and the gradient of mobile phase A was held at 98% for 1 min, 98% to 85% in 2 min, 85% to 50% in 1 min, followed by a linear gradient to 5% A in 1 min, held at 5% A for 3 min. The sample volume injected was 5 μL.

### Feeding experiment of bacterial symbiont *E. casseliflavus*

The healthy third-instar larvae were fed on the antibiotics diet for 2 days to remove the bacterial symbionts (Ant-fed). Then bacterial solution (OD_600_ = 1.0) was washed twice with PBS solution, and resuspended in an equal volume of PBS solution and per-well added 100 μL bacteria-PBS solution. The Ant-fed insects were feeding bacteria-PBS solution for 2 days (BS-fed). Then the insecticide susceptibility and the infection dynamics were assayed (Fig. S5).

### Complete genome sequencing and comparative genomic analysis

The genomic DNA of *E. casseliflavus* was extracted by using the Cetyltrimethyl Ammonium Bromide (CTAB) method with minor modification. The DNA concentration, quality and integrity were determined by using a Qubit Fluorometer (Invitrogen, USA) and a NanoDrop Spectrophotometer (Thermo Scientific, USA). Sequencing libraries were generated using the TruSeq DNA Sample Preparation Kit (Illumina, USA) and the Template Prep Kit (Pacific Biosciences, USA). The genome sequencing was then performed by Personal Biotechnology Company (Shanghai, China) by using the Pacific Biosciences platform and the Illumina Novaseq platform. Data assembly proceeded after removing adapter contamination and filtering data by using AdapterRemoval (Lindgreen, 2012) and SOAPec (Luo et al, 2012). The filtered reads were assembled by Unicycler (Wick et al., 2017). The complete genomic sequence was annotated by eggnog-mapper (http://eggnog-mapper.embl.de/) (Cantalapiedra et al., 2021). The genome sketch was drawn with CGview (https://proksee.ca/) with the clustered regularly interspaced short palindromic repeats (CRISPRs), resistance gene, and ORFs prediction and GS content calculation (Grant and Stothard, 2008).

### Proteome prediction of degradation pathway of chlorantraniliprole

The proteins with chlorantraniliprole-degrading potentially function were screened according to a previous study (Zhang et al., 2022). To mine enzymatic activities and metabolic pathways related to chlorantraniliprole degradation, biodegradation experiments of chlorantraniliprole by *E. casseliflavus* were conducted. Then, both intracellular and extracellular proteins were separately extracted from the cells harvested after 3 days based on the following method: For intracellular proteins extraction, the collected cells of *E. casseliflavus* in two groups (with or without chlorantraniliprole) were lysed by using Branson SFX550 Ultrasonicator, and then the solution was centrifuged to obtain supernatant as the intracellular protein’s solution. For extracellular protein extraction: acetone precipitation method was used to recover extracellular proteins, and the specific steps were described as follows: a mixture of extraction solution and pre-cooled acetone (v:v⎕=[1:1) was stirred for 1[h at 0 °C, and then placed on 4°C overnight. The mixture in each group was concentrated (10,000 rpm, 4°C) to harvest the proteins.

The ability of the protein solutions to degrade chlorantraniliprole was further tested in vitro. The following experiments were performed in a 2-mL system: (1) EC: MSM liquid medium + *E. casseliflavus* (OD_600_[=[0.1) + chlorantraniliprole (20 mg/L); (2) IEC: MSM liquid medium + chlorantraniliprole (20 mg/L) + intracellular protein (100 μg/mL); (3) OEC: MSM liquid medium + chlorantraniliprole (20 mg/L) + extracellular protein (100 μg/mL); (4) Control: MSM liquid medium + chlorantraniliprole (20 mg/L). All treatments were cultured in a shake (150⎕rpm, 30⎕°C) for 3 days and the chlorantraniliprole content was measured via the method described previously (Fig. S6a).

Mass spectrometry analysis of proteomic proteins was resolved with a Thermo Ultimate 3000 integrated nano-HPLC system that directly interfaced with a Thermo Orbitrap Fusion Lumos mass spectrometer (LC-MS/MS) to explore chlorantraniliprole degradation-related proteins. The sodium dodecyl-sulfate polyacrylamide gel electrophoresis (SDS-PAGE) was used to separate the protein extracts and stained with Coomassie Blue G-250. The gel bands of the target were cut into pieces. The sample was digested by trypsin with prior reduction and alkylation in 50⎕mM ammonium bicarbonate at 37⎕°C overnight. The digested products were extracted twice with 1% formic acid in a 50% acetonitrile aqueous solution and dried to reduce volume by speed Vacuum Concentrator. The peptides were separated by a 65⎕min gradient elution at a flow rate of 0.3⎕µL/min with the Thermo Ultimate 3000 integrated nano-HPLC system directly interfaced with the Thermo orbitrap fusion lumos mass spectrometer. The analytical column was a homemade fused silica capillary column (75⎕µm ID, 150 mm length; Upchurch, Oak Harbor, WA) packed with C-18 resin (300⎕A, 3⎕µm, Varian, Lexington, MA). Mobile phase A consisted of 0.1% formic acid, and mobile phase B consisted of 80% acetonitrile and 0.1% formic acid. The mass spectrometer was operated in the data-dependent acquisition mode using the Xcalibur 4.1 software and there is a single full-scan mass spectrum in the Orbitrap (300–1800⎕m/z, 60,000 resolution) followed by 20 data-dependent MS/MS scans at 30% normalized collision energy. Each mass spectrum was analyzed using the Peak studio for database searching. The reference strain is *E. casseliflavus* (strain EC20, NC_020995.1). The protein abundance was then analyzed by R package DEP (version 1.22.0). The Volcanic map and heatmap were drawn with the online software Omicstudio (https://www.omicstudio.cn/index).

### *E. casseliflavus* infection detection and transmission experiment

To detect whether *E. casseliflavus* can infect the host effectively, functional gene that distinguishes *E. casseliflavus* from other closely related bacterial symbionts or host DNA were first screened. To this end, BPGA and anvo’o (Chaudhari et al., 2016; Eren et al., 2021) based pangenomic analysis was used to identify the unique protein cluster of *E. casseliflavus* compared with other *Enterococcus* bacteria strains (including 2 *E. casseliflavus*: NC_020995.1 and NZ_CP086066.1, an *Enterococcus durans*: NZ_CP022930.1; and an *Enterococcus mundtii*: NZ_CP018061.1). A hypothetical protein of *E. casseliflavus* (EMBL3) was used to design qRT-PCR primer the following: ECHP F (5’-GTTCCCTGGTGCTTATGGCT-3’) and ECHP[R (5’-AATTCTGCATCGCAGGGGAA -3’) by NCBI primer BLAST. The primers availability was then verified in a series of concentrations of templates: with or without *E. casseliflavus* infection hosts DNA and cDNA. To detect the infection rate of *E. casseliflavus* in *S. frugiperda*, insect’s individual was used for DNA extraction and each field population performed 30 insects. The infection rate was detected via PCR by using a specific primer of *E. casseliflavus* above.

To explore the transmission pattern of *E. casseliflavus* between individuals and a population (Nanning) of *S. frugiperda*, *E. casseliflavus-*free *S. frugiperda* were fed on the artificial diet containing *E. casseliflavus* for 2 days, then put them in normally artificial diet, the insects were sampled and *E. casseliflavus* infection detected throughout their development up to the larvae of the next generation. For horizontal transmission, a diet crossover and fratricide experiment were conducted separately. In brief, a 5^th^-instar *E. casseliflavus* free *S. frugiperda* and a 3^rd^-instar *E. casseliflavus* infected *S. frugiperda* were reared together with or without an artificial diet. Different insect instar representing different insect sizes can distinguish whether the insect is a *E. casseliflavus* free or *E. casseliflavus* infected state at the beginning of the experiment. After 2 days, each *E. casseliflavus* free *S. frugiperda* was collected and detected *E. casseliflavus* infection.

### Data bioinformatics, statistics, and visualization

The LC_50_ of *S. frugiperda* to each insecticide was calculated by Polo Plus (ProbitLogitAnalysis, Le-OraSoftware) and the survival curve analysis was performed by GraphPad Prism (Version 6.0). The 16S rDNA gene amplicon sequence data were analyzed by QIIME2 with demultiplexed using the demux plugin followed by primer cutting with the cutadapt plugin. Sequence data were processed with the DADA2 plugin to quality filter, denoise, merge and remove chimeras. Non-singleton amplicon sequence variants (ASVs) were aligned with maft. Taxonomy was assigned to the ASVs using the classify-sklearn naïve Bayes taxonomy classifier in the feature-classifier plugin, against the Greengenes Database (http://greengenes.lbl.gov). The α-diversity and β-diversity were estimated using the diversity plugin with rarefied samples, including four α-diversity indices [Shannon diversity index and observed ASVs]. The VEGAN package (version 2.5-7) was used to calculate the Bray–Curtis distances between samples with the ASVs table, and principal coordinate analysis (PCoA) was performed by distance matrices using the cmdscale function in R. All results from R were visualized and plotted using ggplot2 (3.3.3). GraphPad Prism version 6.0 and Omicstudio were used for statistical analyses and data plotting. Statistical comparisons were performed with Student’s *t*-test, and the survival curve was analyzed by the log-rank (Mantel–Cox) test. The significance level was set at \**P*⎕<⎕0.05 (and different letters) and \*\**P*⎕<⎕0.01.

## Data Availability

The sequencing raw data generated in this study are deposited in the China National GeneBank DataBase (CNP0004265), and the mass spectrometry proteomics data have been deposited to the ProteomeXchange Consortium via the PRIDE partner repository with the dataset identifier PXD045281.

## Supporting information

Supplemental table and fugue

## Acknowledgement

This work was supported by the HRHI program 202309010 of Westlake Laboratory of Life Sciences and Biomedicine, the China Postdoctoral Science Foundation (Certificate Number: 2023M733188), the National Natural Science Foundation of China under Grant No. 32302393, the Research Center for Industries of the Future (Grant No. WU2022C030) and the Westlake Center for Synthetic Biology and Integrated Bioengineering (WU2022A008) at Westlake University. The authors thank Ms. Yisong Xu for laboratory management support and also thank Dr. Zhe Zhang for bacterial isolates support. The authors thank the Westlake University HPC Center for computation support.

