## Supplemental table and fugue for "Uninheritable but widespread bacterial symbiont mediates insecticide detoxification of an agricultural invasive pest *Spodoptera frugiperda*"

Yunhua Zhang and Feng Ju*

**Figure and Table Legends**

**Table S1 The influence of *E. casseliflavus* on the susceptibility of *Spodoptera frugiperda* to chlorantraniliprole.**

**Table S2 KEGG enrichment of difference expression extracellular and intracellular protein in *E. casseliflavus* exposure with or without chlorantraniliprole.**

**Table S3 *Spodoptera frugiperda* field population information used in this study.**

**Table S4 Detailed information of insecticides used in this study.**

**Fig. S1 The influence of antibiotic on the bacterial symbionts of *Spodoptera frugiperda.***

**Fig. S2** **The influence of insecticide exposure on the microbiome of *Spodoptera frugiperda.***

**Fig. S3 Full-length 16S rRNA gene-based phylogenetic tree of four bacterial symbionts isolate from *Spodoptera frugiperda***

**Fig. S4 Degradation of *Enterococcus casseliflavus* on chlorantraniliprole.**

**Fig. S5 Experiment design of the influence of *Enterococcus casseliflavus* feeding on the susceptibility of *Spodoptera frugiperda* to chlorantraniliprole.**

**Fig. S6 Biodegradation potential and proteome analysis of intracellular and extracellular proteins of *Enterococcus casseliflavus* to chlorantraniliprole.**

**Fig. S7 Molecular signal of potential chlorantraniliprole-biodegrading products in *Enterococcus casseliflavus* treated chlorantraniliprole solution.**

**Fig. S8 Biodegradation pathway of chlorantraniliprole prediction with EAWAG-PPS.**

**Fig. S9 Genomic map of *Enterococcus casseliflavus***

### Supplementary Tables

**Table S1 The influence of *E. casseliflavus* on the susceptibility of *Spodoptera frugiperda* to chlorantraniliprole.**

| **Treatment** | **N** | **Slope (SE)** | **LC_50_ (95%CI)** | **ꭕ^2^ (df)** |
| --- | --- | --- | --- | --- |
| Control | 144 | 1.41 (0.28) | 14.80 (8.95-36.56) | 3.83 (3) |
| BS-fed  (*E. casseliflavus*) | 144 | 1.66 (0.28) | 15.33 (10.33-29.30) | 2.09 (4) |
| Ant-fed | 144 | 1.55 (0.26) | 3.23 (2.00-4.59) | 0.71 (4) |

**Table S2 KEGG enrichment of difference expression protein in IE vs. IEC and OE vs. OEC.**

| **Treatment group** | **Pathway name** | **ID** | **Input number** | **Background number** | ***P* value** | **Protein (Uniprot ID)** |
| --- | --- | --- | --- | --- | --- | --- |
| IE vs. IEC | Aminoacyl-tRNA biosynthesis | ecas00970 | 4 | 27 | 0.0030 | C9A7W7, C9A757, C9A7J2 and C9ABG4 |
|  | Ascorbate and aldarate metabolism | ecas00053 | 2 | 9 | 0.019 | C9A681 and C9A9X1 |
|  | Bacterial secretion system | ecas03070 | 2 | 10 | 0.022 | C9A4U7 and C9A8R5 |
|  | Protein export | ecas03060 | 2 | 18 | 0.058 | C9A4U7 and C9A8R5 |
| OE vs. OEC | Metabolic pathways | ecas01100 | 130 | 540 | 7.90E-05 | C9AB43, C9A551, C9A559, C9A558, C9AC27, C9AA22, C9A8B5, C9A8B3, C9AD80, C9A5N7, C9AD77, C9ACV8, C9A893, C9A890, C9A4D4, C9A4D7, C9A5G2, C9A715, C9A7X3, C9A7X0, C9ABN0, C9A7W4, C9A6Q9, C9A7W1, C9A7W8, C9ACF5, C9A8J3, C9A7H5, C9AB99, C9A9U5, M9T8C8, C9ABM9, C9AB86, C9A8U3, C9AD27, C9A4W5, C9A9D0, C9A9D1, C9A9D5, C9ABC7, C9ACA6, C9A582, C9AB95, C9AC58, C9A502, C9AB19, C9A7B2, C9A9E7, C9A9E3, C9ABL3, C9A6F1, C9ACQ0, C9A6W9, C9ACH4, C9ACH5, C9ABS8, C9A9Z5, C9A8F3, C9ACB1, C9A996, C9AC63, C9AD78, C9ADC8, C9A4Y3, C9A7K7, C9A4N2, C9AD51, C9A4N8, C9A6M7, C9A4Q3, C9ACI3, C9A8A5, C9A9P8, C9A8A9, C9A9C1, C9A7S9, C9A7S8, C9AAJ4, C9A9X3, C9ABV4, C9AAS8, C9A6L7, C9A4P0, C9ACS5, C9AAS2, C9ACS1, C9A598, C9A8H0, C9ABX7, C9A4P1, C9AAX8, C9A4K2, C9A762, C9A763, C9AB65, C9A4H1, C9AAK3, C9A4S0, C9AAP1, C9AAK9, C9A7E1, C9A7M5, C9A7M3, C9ABP8, C9A7N4, C9A7T2, C9A7T0, C9AD09, C9ABG8, C9ABG9, C9AC98, C9A9N1, C9A8C0, C9A8C3, C9AC10, C9AAK5, C9A8X6, C9AC18, C9AAK4, C9A5I7, C9A8P6, C9A5I2, C9A4R9, C9A5I9, C9A5H8, C9A7U9, C9A7N7, C9A7N8, C9A7U1 and C9A4C9 |
|  | Biosynthesis of secondary metabolites | ecas01110 | 57 | 205 | 0.00071 | C9AB43, C9A8F3, C9A9E7, C9A996, C9AC63, C9AD78, C9A5G2, C9ADC8, C9A8H0, C9A4P1, C9A9U5, C9A890, C9A762, C9A763, C9AD80, C9A7X0, C9ABM9, C9AD77, C9ACX2, C9A4N8, C9A5H8, C9A6M7, C9A4Q3, C9A893, C9ACI3, C9ABC7, C9A4D7, C9A715, C9AC98, C9A8Z1, M9T8C8, C9A582, C9ABP8, C9A6L7, C9A9P8, C9A8X6, C9AD51, C9AC18, C9A5I7, C9ABN0, C9A7S9, C9A8P6, C9A9E3, C9AAJ4, C9AC58, C9AAS2, C9AAS8, C9A7N4, C9A7T0, C9A7N7, C9A7N8, C9A4P0, C9ACS5, C9ACH5, C9A4C9, C9ACS1 and C9A7H5 |
|  | Ribosome | ecas03010 | 20 | 57 | 0.0056 | C9ACL2, C9ABD5, C9A9G6, C9AAP2, C9ABC0, C9A9M7, C9AB49, C9A9H6, C9A7A2, C9A9G3, C9A9F8, C9A9H2, C9A9G5, C9A7I0, C9A9G7, C9A4V4, C9A9G1, C9A4V2, C9AAR4 and C9A9H1 |
|  | Cysteine and methionine metabolism | ecas00270 | 11 | 28 | 0.020 | C9A8F3, C9A7S9, C9AAJ4, C9ABL3, C9ADC8, C9A5I9, C9AAS8, C9A6M7, C9A7T2, C9A7T0 and C9A7H5 |
|  | Propanoate metabolism | ecas00640 | 8 | 18 | 0.027 | M9T8C8, C9A7S8, C9A4N2, C9A9N1, C9ADC8, C9A7N4, C9A7N7 and C9A7N8 |
|  | Aminoacyl-tRNA biosynthesis | ecas00970 | 10 | 27 | 0.034 | C9ACW2, C9A4H2, C9A4D0, C9A7Q7, C9A8U3, C9ABG4, C9A7W8, C9AB86, C9ACG4 and C9A8U5 |
|  | Glycerophospholipid metabolism | ecas00564 | 6 | 13 | 0.047 | C9A4D7, C9A581, C9A7X0, C9A8C3, C9ACQ3 and C9A4Q3 |

Table S3 ***Spodoptera frugiperda* field population information used in this study.**

| **Population** | **Host plant** | **Province, city** | **Site** | **Date** |
| --- | --- | --- | --- | --- |
| AH | Corn | Anhui, Wuwei | 117.91 °E，31.15 °N | 2022-08-31 |
| GX | Corn | Guangxi, Nanning | 108.25 °E, 22.85 °N | 2022-09-16 |
| GD | Corn | Guangdong, Zhanjiang | 110.23 °E, 20.73 °N | 2022-08-17 |
| HB | Corn | Hubei, Jingzhou | 112.14 °E, 30.20 °N | 2022-07-20 |
| ZJ | Sorghum | Zhejiang, Dongyang | 120.32 °E, 29.28 °N | 2022-08-08 |

Table S4 Detailed information of insecticides used in this study.

| Insecticides | Purity | Types | Producers |
| --- | --- | --- | --- |
| Indoxacarb | 95% | Carbamic esters | HuBei WeiDeLi Chemical Technology Co., Ltd |
| Emamectin benzoate | 95% | Avermectins | InNer MonGoLia Jumbo Biochemistry Co., Ltd |
| Lufenuron | 97% | Benzoylureas | JiangSu FengShan Group Co., Ltd |
| Lambda-cyhalothrin | 96% | Pyrethroids | ShanDong WeiFang Rainbow Chemical Co., Ltd |
| Chlorantraniliprole | 98% | Diamides |  |
| Chlorfenapyr | 97% | Pyrroles |  |

### Supplementary Figures

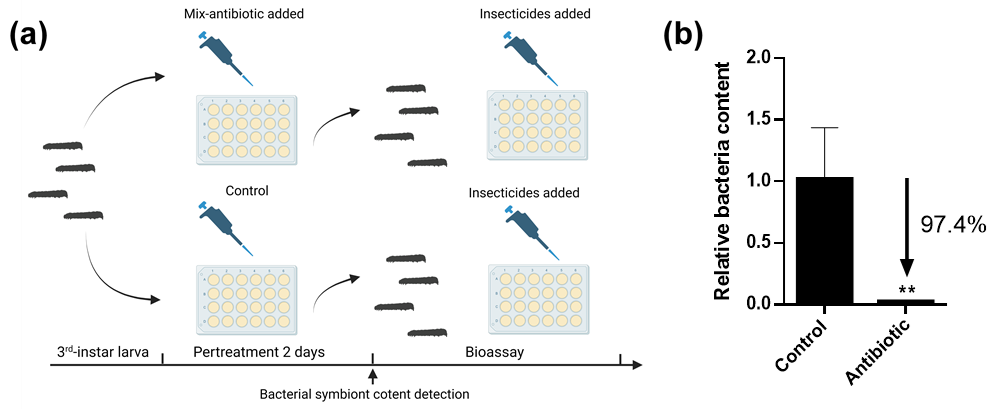

**Fig. S1 The influence of antibiotic on the bacterial symbionts of *Spodoptera frugiperda.***

Note: Bar-plot shown SEM and “**” represent there is a significantly difference between control and antibiotic treatment (*P* < 0.01 by student’s *t* test).

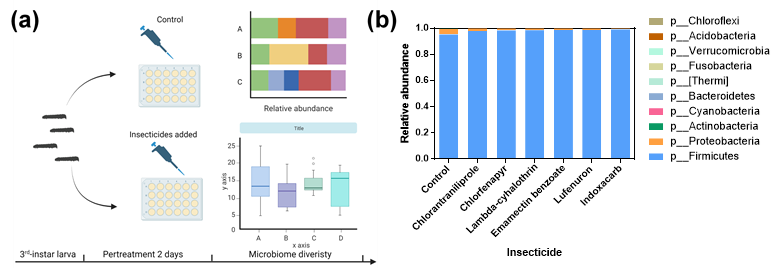

**Fig. S2** **The influence of insecticide exposure on the** **microbiome of *Spodoptera frugiperda.***

**(a)**: Experiment design; **(b)**: Microbiome structure at phylum level of *S. frugiperda* in response to insecticide exposure.

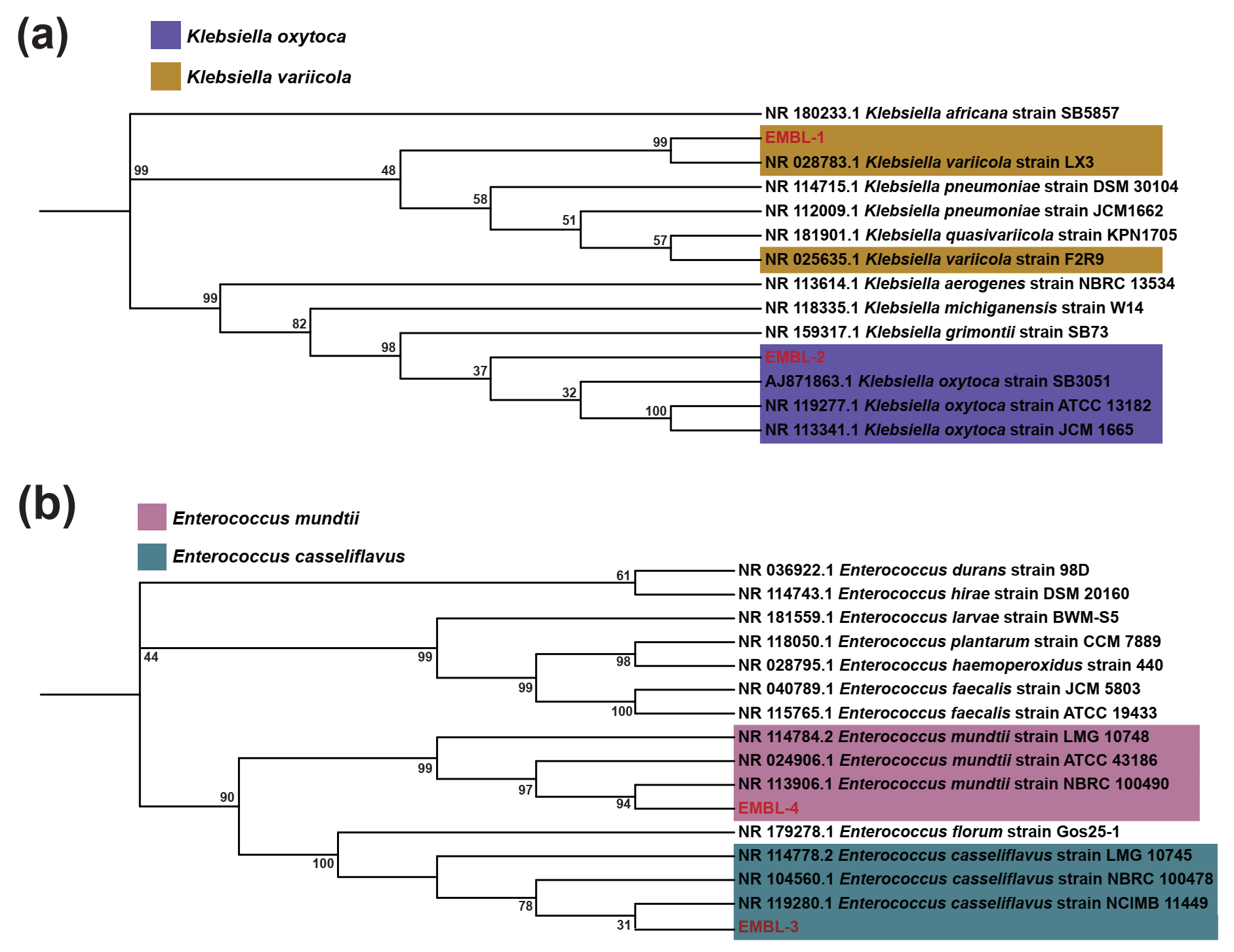

**Fig. S3 Full-length 16S rRNA gene-based phylogenetic tree of four bacterial symbionts isolate from *Spodoptera frugiperda*. (a):** *Klebsiella*; **(b):** *Enterococcus.*

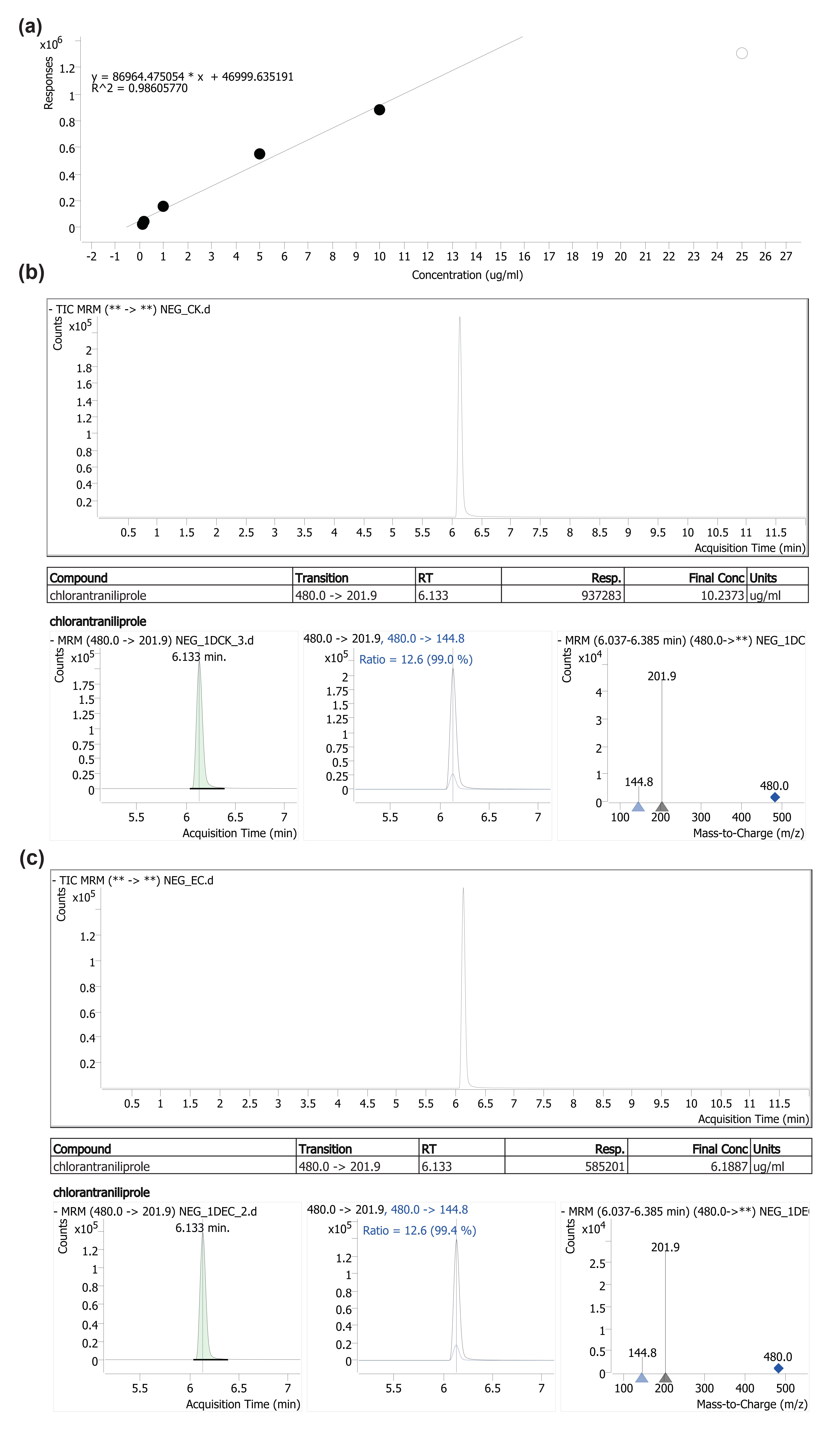

**Fig. S4 Degradation of *Enterococcus casseliflavus* on chlorantraniliprole. (a):** Standard curve; **(b and c):** The concentration of chlorantraniliprole in control **(b)** and *Enterococcus casseliflavus* treatment **(c)** group.

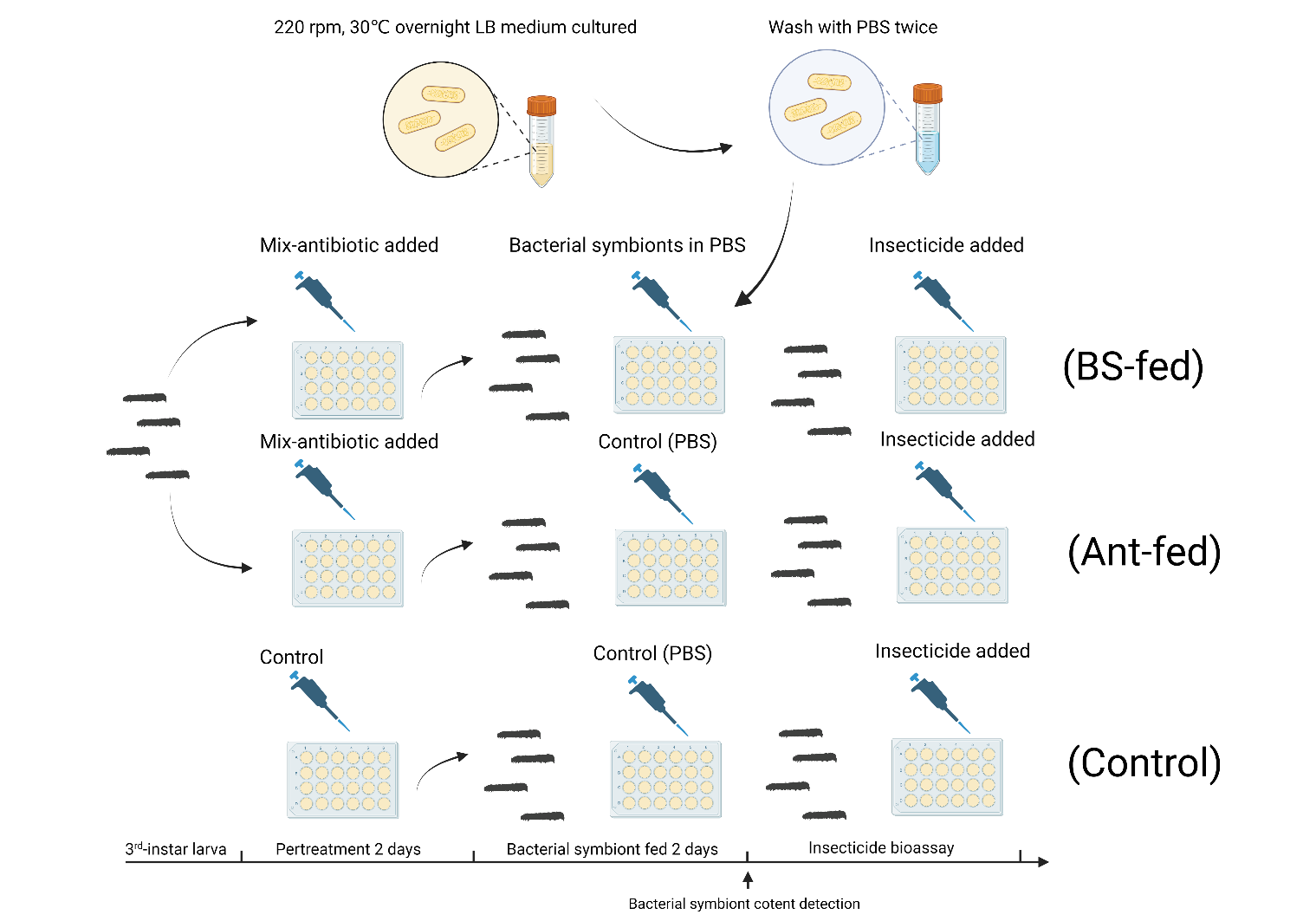

**Fig. S5 Experiment design of the influence of *Enterococcus casseliflavus* feeding on the susceptibility of *Spodoptera frugiperda* to chlorantraniliprole.**

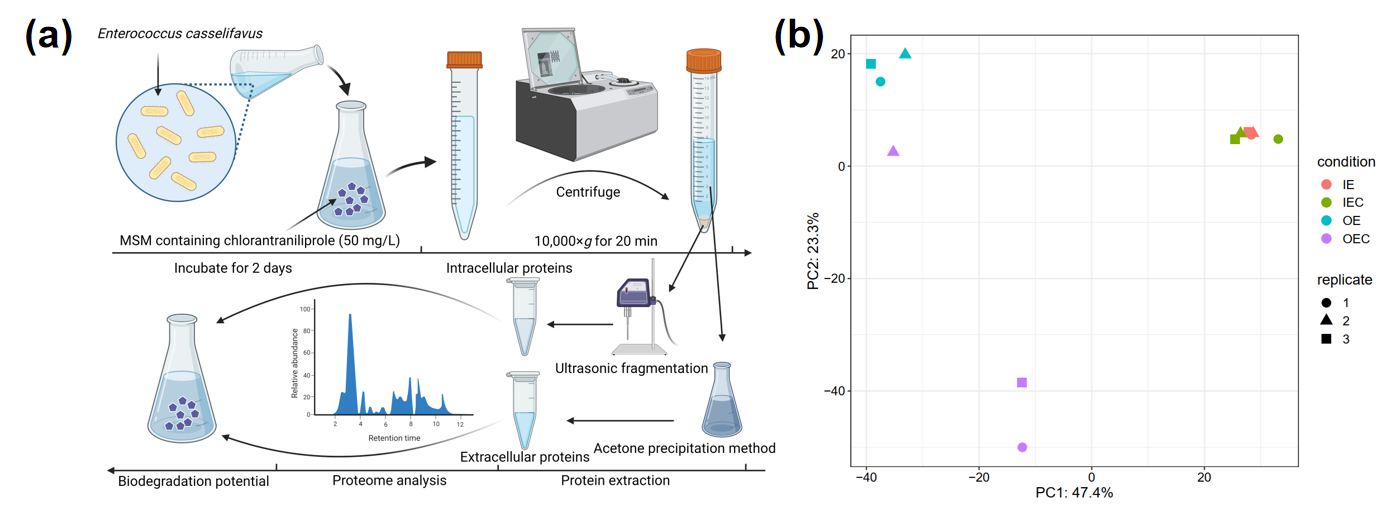

**Fig. S6 Biodegradation potential and proteome analysis of intracellular and extracellular proteins of *Enterococcus casseliflavus* to chlorantraniliprole. (a):** Experiment design; **(b):** PCA analysis of intracellular and extracellular proteins of *Enterococcus casseliflavus* with or without chlorantraniliprole exposure. OE: Extracellular proteins expression level in MSM + *E. casselifiavus*; OEC: Extracellular proteins expression level in MSM + *E. casselifiavus* + chlorantraniliprole; IE: Intracellular proteins expression level in MSM + *E. casselifiavus*; IEC: Intracellular proteins expression level in MSM + *E. casselifiavus* + chlorantraniliprole.

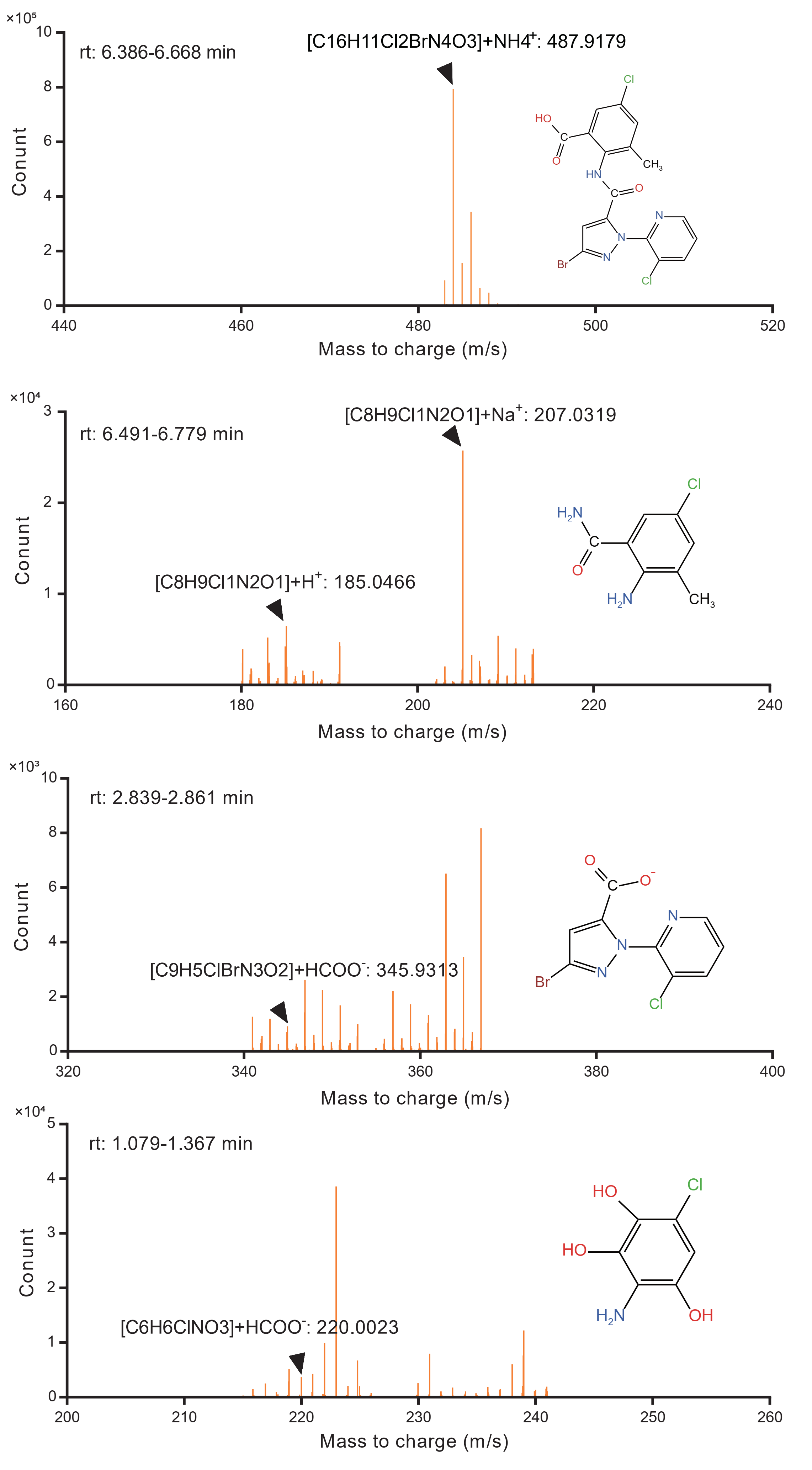

**Fig. S7 Molecular signal of potential chlorantraniliprole-biodegrading products in *Enterococcus casseliflavus* treated chlorantraniliprole solution.**

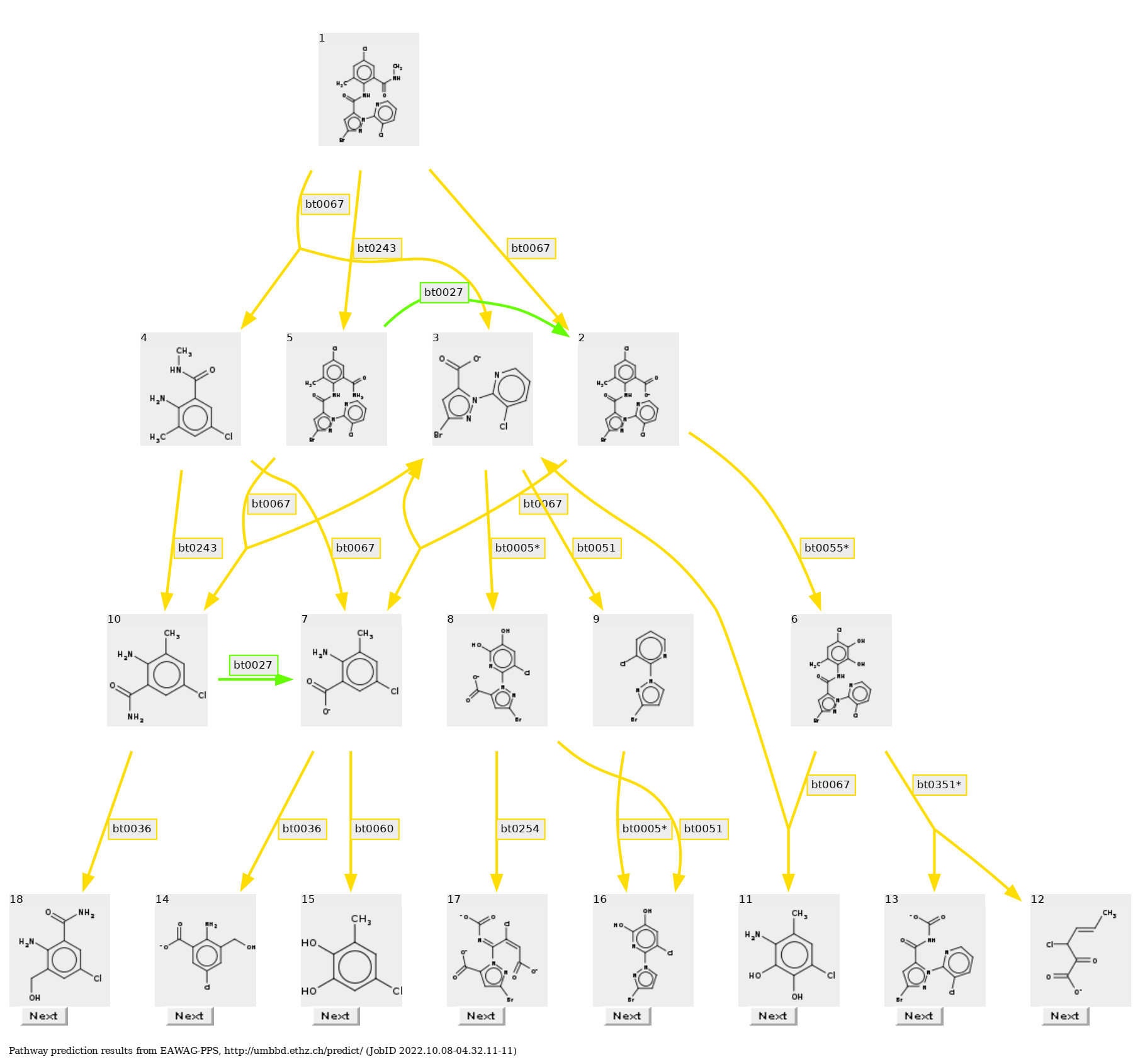

**Fig. S8 Biodegradation pathway of chlorantraniliprole prediction with EAWAG-PPS.**

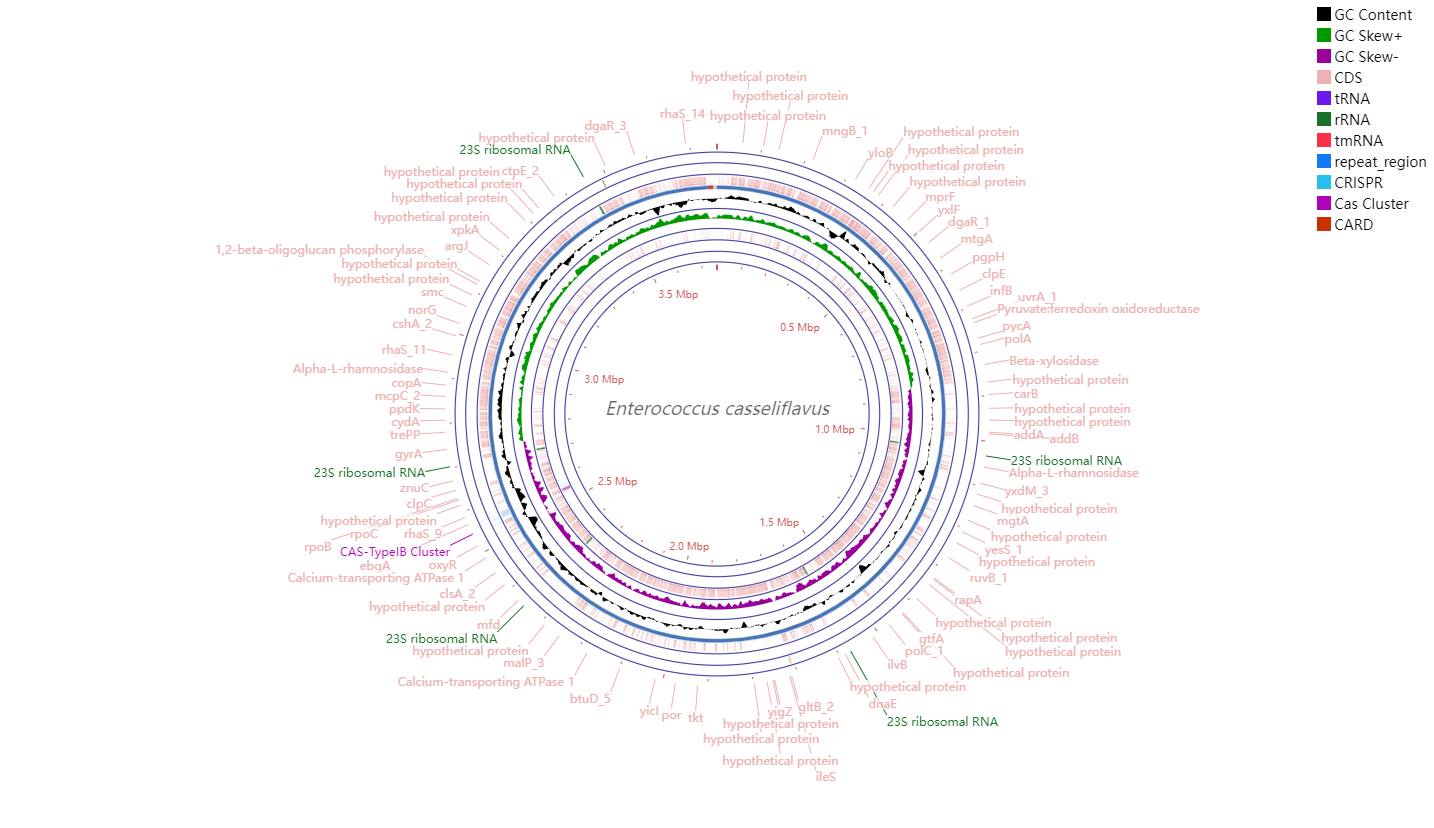

**Fig. S9 Genomic map of *Enterococcus casseliflavus* EMBL-3**
